# A novel link between Cell Wall Integrity pathway and Flocculation, regulated by yeast Sen1 dependent interplay between Rlm1 and Tup1

**DOI:** 10.1101/463703

**Authors:** Santhosh Kumar Sariki, Ramesh Kumawat, Vikash Singh, Raghuvir Singh Tomar

## Abstract

The budding yeast, *Saccharomyces cerevisiae* is one of the most studied organisms used for the synthesis of products, to explore the human diseases and eukaryotic gene expression mechanisms. The yeast cells with flocculation property are in high demand for industrial applications. However, the pathogenic yeast becomes drug-resistant due to flocculation/biofilm phenotype. The flocculation property of yeast depends on the expression of specific *FLO* genes. Genetic and epigenetic factors have been suggested to induce the expression of *FLO*s and flocculation, an evolutionarily conserved process. The present study was undertaken to identify a molecular link between stress caused by genetic and epigenetic factors and expression of *FLO*s. We utilized flocculating yeast strains to study the regulation of *FLO* genes and flocculation phenotype. We found rough surface morphology and constitutive activation of Slt2 in flocculating cells. The external cell wall stress factors as well as specific mutations in Sen1 and histone proteins strongly correlated with the induction of *FLO* genes whereas deletion of *SLT2*/*RLM1*, suppressed the expression and flocculation phenotype. We detected constitutive binding of Rlm1 and eviction of Tup1 from the promoters of *FLO1* and *FLO5* genes in flocculating cells. Thus we provide evidence for the CWI pathway dependent flocculation of yeast, regulated by Sen1 mediated interplay between Tup1 and Rlm1.

## 1. Introduction

Gene expression mechanisms in eukaryotes are tightly regulated by complex interplay between regulatory factors (Smith and Shilatifard 2010; Smith and Workman 2012). Often regulatory factors compete for their binding to genes. The cooperation among the elements of transcriptional machinery maintains cellular homeostasis. For survival, it is essential to generate an appropriate transcriptional outcome in response to extrinsic and intrinsic factors (Mcmurray and Tainer 2003; Thorsen *et al.* 2006; Vihervaara *et al.* 2018). Living organisms are constantly exposed to a wide variety of agents which can induce genetic and epigenetic changes (Baccarelli and Bollati 2009). Mutations in activators or repressors of transcription apparatus have been linked with a variety of diseases; including cancer and developmental disorders (Lee and Young 2013). The survival in response to such perturbations is dependent on coordination among factors of specific molecular signaling pathways and transcriptional mechanisms (Taymaz-nikerel *et al.* 2016). For example, upon exposure to the cell wall (Garcia *et al.* 2004; Levin 2005) and DNA damaging agents (O’Connor 2015), specific cell wall sensor proteins transmit the signal to nucleus through ‘Cell Wall Integrity’ (CWI) (Jung and Levin 1999) and genomic integrity signaling pathways which subsequently triggers the de-repression of cell wall and DNA repair genes respectively. In the absence of DNA damage, transcription of repair genes is turned off by the recruitment of repressor complex leading to the formation of transcription resistant chromatin structure (Smith and Johnson 2000).

Yeast mitogen-activated protein kinase (MAPK) signaling pathways transduce external signals via Slt2/Mpk1 and Hog1 kinases, generates appropriate cellular responses (Chen and Thorner 2007; Gustin *et al.* 1998) which are required for survival under cell wall and osmotic stress respectively (Bermejo *et al.* 2008; De Nobel *et al.* 2000; Levin 2011). Zymolyase treatment induces the phosphorylation of Slt2 and Hog1 to activate the expression of the CWI pathway genes (Garcia *et al.* 2009). Under cell wall stress conditions, sensor proteins activate a kinase cascade; Rom2-Rho1-Pkc1 which integrates into the MAP Kinase module which composed of Bck1, Mkk1, and Mkk2 proteins. The active form of Bck1 activates two kinases; Mkk1 and Mkk2, which further phosphorylates a downstream MAPK, Slt2 (Kim *et al.* 2010). The phosphorylated form of Slt2 subsequently enters the nucleus and activates two transcription factors; Rlm1 and SBF (Garcia *et al.* 2016; Madden *et al.* 1997). For example; the transcriptional induction of one of the CWI genes occurs in response to cell wall damage through the Slt2 activation-dependent recruitment of Rlm1, SWI2/SNF2 and SAGA complexes (Sanz *et al.* 2012; Sanz *et al.* 2016). Under non-inducing conditions, transcription of CWI genes is terminated by the binding of Nrd1-Nab3-Sen1 (NNS) complex. However, under cell wall stress conditions, Mpk1 mediates the up-regulation of CWI genes by blocking recruitment of NNS transcription termination complex (Kim and Levin 2011). Furthermore, the significance of MAPK pathway has also been implicated in the filamentous growth of pathogenic yeast due to up-regulation of cell-cell adhesion/flocculation genes (Chavel *et al.* 2014; Chavel *et al.* 2010). Although above studies suggest that the expression of CWI and flocculation genes are probably regulated through a common MAP kinase signaling dependent mechanism, however, a molecular link between these two is yet to be established.

The genetic, epigenetic and environmental factors have been shown to induce flocculation phenotypes (Kim and Rose 2015; Liu *et al.* 2015; Soares 2011) such as mutations in certain genes, nutritional stress, temperature, pH, oxygen supply and alteration in cell wall composition. The flocculation or aggregation of cells is also a property of yeast and bacteria to form biofilm and become resistant to drugs (Chandra *et al.* 2001). Yeast also forms multi-cellular flocs/clumps under adverse growth or particular physiological conditions such as lipid unsaturation and limited oxygen supply (Degreif *et al.* 2017). Induction of flocculation protects cells from such stressful environmental situations and allows them to survive for long (Hope *et al.* 2017). There are six major *FLO* genes in yeast; *FLO1, FLO5, FLO8, FLO9, FLO10*, and *FLO11*, which are responsible for flocculation and biofilm formation. The role of Flo1 is quite well established in flocculation, bio-film like phenotype of yeast (Goossens *et al.* 2011; Sim *et al.* 2013; Smukalla *et al.* 2008). The Flo8p is considered as a transcription factor to drive the expression of *FLO11* (Fichtner *et al.* 2007), and significance of *FLO11* has been implicated in the mating process of yeast (Guo *et al.* 2000) and invasive growth.

A transcription factor, Mss11 is also known to regulate the expression of *FLO10* and *FLO11* (Bester et al. 2012) essential for cellular adhesion phenotype. The expression of *FLO11* is linked with mutations in certain genes leading to flocculation of yeast. The *FLO11*-associated phenotypes such as biofilm formation and adhesion to polystyrene are inhibited by L-Histidine through altering the chitin and glycan content on the cell wall of flor yeasts (BOU ZEIDAN *et al.* 2014). Similarly, the deletion of the ribosomal *RPL32* gene in fission yeast (Liu *et al.* 2015) and deletion mutants of the COMPASS complex (Histone H3 methyltransferase) in budding yeast induces the flocculation phenotype (Dietvorst and Brandt 2008).

Sen1p is a member of Nrd1-Nab3-Sen1 (NNS) complex, primarily involved in termination and processing of non-coding transcripts through the process of nuclear exosome complex mediated degradation of RNA (Chinchilla *et al.* 2012; Finkel *et al.* 2010). Role of Sen1p has also been implicated in genomic integrity by preventing co-transcriptional R-loop formation (Cohen *et al.* 2018; Grunseich *et al.* 2018). Defects in Sen1p are associated with down-regulation of DNA repair genes and sensitivity to DNA damaging agents (Golla *et al.* 2013). Furthermore, significant alteration in the expression of lipid homeostasis pathway genes is reported (Sariki *et al.* 2016) in *Sen1* mutants. Moreover, many mutations in Senataxin, the human homolog of yeast Sen1 are correlated with genetic disorders, ALS4, and AOA2 (Groh *et al.* 2017). However, the underlying mechanism is yet to be elucidated for the development of disease pathogenesis. All the fungal cell wall adhesins/flocculins, as well as membrane adhesion proteins of higher eukaryotes, are Gpi-anchored (Saha *et al.* 2016; Verstrepen and Klis 2006). In fungus, these glycosyl-phosphatidylinositol (GPI) anchored glycoproteins are essential for invasive growth, cell-cell adhesion, and mating (Guo *et al.* 2000). The conserved amyloid-like amino acid sequences are also found in yeast cell adhesins; Flo1p and Flo11p (Lipke *et al.* 2012; Rameau *et al.* 2016; Ramsook *et al.* 2010) which suggest that cellular aggregation by amyloid sequence containing proteins is the probable reason for neurological diseases in humans, and their localization on the yeast cell wall contributes in cell-cell aggregation, leading to biofilm formation.

Recently, upregulation of Flo1p has been observed in *Rpb11* and *Sen1* mutant yeast cells (Chen *et al.* 2017) supporting our earlier observations. However, the molecular mechanism has not been explored. The deletion of *TUP1* in budding yeast induces strong flocculation phenotype (Lipke and Hull-Pillsbury 1984). The yeast *FLO1* and DNA repair genes are repressed by a global repressor Cyc-Tup1 complex along with histone deacetylases; Hda1 and Rpd3 (Church *et al.* 2017; Zhang and Reese 2004). During DNA damaging conditions or in the absence of Cyc-Tup1 repressor, the occupancy of Hda1 and Rpd3 decreases whereas occupancy of Swi/Snf increases (Fleming *et al.* 2014; Fleming and Pennings 2001). Thus the genes of DNA repair and flocculation are regulated by the antagonistic actions of chromatin remodeling and repressor complexes (Fleming and Pennings 2001).

We have earlier reported strong flocculation phenotype of a few *Sen1* mutant strains of *Saccharomyces cerevisiae* (Singh *et al.* 2015) as well as down-regulation of *SOD1* (Sariki *et al.* 2016). Another study has shown higher chitin content in the cell wall of *sod1∆* cells (Liu *et al.* 2010) indicating the role of Sen1 in cell wall maintenance. However, why do mutations in *Sen1* result in flocculation phenotype is not clear. In the present study through our extensive investigations, we have established a Sen1 regulated novel link between the CWI pathway and flocculation. We utilized flocculating mutant strains of *Sen1* and histones as well as *tup1∆* cells for the studies. We found that flocculating cells exhibit rough surface morphology, enlarged cell size, higher cell wall chitin content, sensitivity to cell wall perturbing agents and constitutive activation (phosphorylation) of Slt2. Our careful analysis of upstream promoter sequences of *FLO* genes suggested a binding site for Rlm1. Indeed through chromatin immunoprecipitation assays; we revealed constitutive binding of Rlm1, TBP, Pol-II and the loss of Tup1 occupancy at the *FLO1* and *5* genes in flocculating *Sen1* mutant cells. The growth of flocculating cells was impaired in the presence of cell wall damaging agents probably due to alteration in cell wall chitin content. The phosphorylation of Slt2 and de-repression of *FLO* genes was also found in wild-type strain in response to stress conditions such as temperature and cell wall damaging agent. Moreover, the drastic decrease in basal level expression of *FLO1, 5, 9, 10* genes in deletion mutants of CWI pathway was observed which further suggest that CWI pathway is required for the regulation of *FLO* genes and flocculation phenotype. The mutations in Sen1p and histones, and exposure to external stress factors, resulted in transcriptional induction of *FLO*s. Interestingly the expression of *FLO* genes was significantly reduced after the deletion of *SLT2* and *RLM1*, leading to suppression of flocculation phenotype. Thus we provide evidence that the CWI pathway plays an essential role in the regulation of yeast flocculation. Altogether for the first time, we established the role of ‘Cell Wall Integrity’ signaling pathway in yeast Flocculation, regulated by Sen1 mediated interplay between Tup1 and Rlm1.

## 2. Materials and Methods

### 2.1. Strains, chemicals, growth media and conditions

The *Saccharomyces cerevisiae* strains used in this study were in the BY4741 background and had been listed in Table S1. The library of histone mutants was purchased from Dharmacon, Cat# YSC5106. The diploid gene deletion library mutants were purchased from Open Biosystems, Cat# YSC1056. Gene tagging and deletions were performed by polymerase chain reaction (PCR) based methods (Janke *et al.* 2004; Wach 1996) using primers listed in Table S2. For the selection of positive colonies, transformants were plated on YPD medium containing G418 sulfate (300μg/ml), and the deletions were verified by PCR using ORF-specific primers. All gene deletions and tagging were confirmed by either PCR or western blotting. *SLT2* deletion was confirmed by western blotting using Anti-Mpk1 antibody and PCR with *SLT2* ORF primer. *RLM1* deletion was confirmed by PCR using *RLM1* primer. Western blot was conducted using an Anti-Myc antibody to verify the genomic Myc tagged Rlm1.

Chemicals used in the experiments were mostly purchased from Sigma Aldrich unless otherwise mentioned. Yeast strains were grown in SC (synthetic complete) media. For making SC media, all amino acids, yeast nitrogen base (YNB), ammonium sulfate and glucose were mixed as per the standard laboratory protocol (Amberg *et al.* 2005). All Yeast strains used in this study were grown at 30°C unless otherwise stated.

### 2.2. Growth sensitivity Assay

Spot assays were performed to investigate the effect of cell wall perturbing agents on the growth of WT and mutants. Mid-log phase cultures of wild-type and mutant yeast cells were 10 fold serially diluted. 3 μl of serially diluted cultures were spotted onto solid Synthetic Complete (SC) Agar plates with or without cell wall perturbing agents; Calcofluor White (CFW), Congo Red (CR) and Caffeine. All plates were incubated at 30°C and growth of the yeast strains was recorded at 72 h by scanning the plates using an HP scanner. For growth curve analysis, exponentially growing cells were seeded in a 96-well cell culture plate in triplicates and incubated in liquid growth media at 30°C with different concentrations of CFW. The OD_600_ at regular time intervals using an automatic plate reader (Eon Microplate

Spectrophotometer) was recorded, and the data were analyzed using GraphPad Prism 5.0 software.

### 2.3. Colony Forming Units

The growth of mutant strains of NNS, TRAMP/Exosome and Histones were determined by CFU assay in the presence of Cell Wall damaging agents was measured by counting CFUs and plotted as plating efficiencies. Saturated cultures of the above-mentioned cells (mutants and respective wild-type cells) were serially diluted, and aliquots containing 1,000 cells were plated on SC Agar plates with and without varying concentrations of Cell Wall damaging agents; CFW (50μg/ml), CR (10μg/ml) and Caffeine (6mM) and incubated at 30°C. Percent cellular viability was determined as the portion of cells that formed viable colonies after 3 days of incubation. This assay was conducted three times. The plating efficiencies were calculated as % of colonies viable from all plated cells.

### 2.4. Microscopy

Images of cells were captured by using an Apotome Carl Zeiss fluorescence microscope through Apochromat 63× objective lens. For the analysis of the cell size of wild-type and mutant strains, cells were grown at 30°C in synthetic complete liquid medium till exponential phase, harvested, washed twice with PBS, visualized and images were acquired using DIC filter.

### 2.5. Flow Cytometry

The intracellular chitin levels and cell size were measured using BD-FACS Aria III equipped with CELL QUEST software. To measure chitin content, WT and mutant strains were grown at 30°C in SC medium till exponential phase and subsequently stained with 50μg/ml CFW for 30 min at 30°C and thereafter cells were collected by centrifugation, washed twice with PBS, re-suspended again in PBS, and fluorescence intensity was measured using DAPI filter. The relative cell size was estimated by plotting the areas of forward (FSC) versus side (SSC) scattering signal parameters.

### 2.6. Scanning Electron Microscopy (SEM)

The SEM analysis of WT and mutants (*sen1-1*, *sen1 ∆N*, and *sen1-K128E*) by following a standard protocol was performed with slight modifications (Tang *et al.* 2014). In brief, 1 ml of overnight cultures were collected by centrifugation and washed twice with dH_2_O before prefixing the cells with 4% Paraformaldehyde (PFA) for 5 min at RT. After PFA fixation, cells were washed twice with 1X PBS and dehydrated serially in different concentrations of ethanol ranging from 30% - 100%, for 10 min in each at room temperature. After final resuspension in 100% ethanol, the cells were sonicated to remove the clumps. Subsequently, cells were diluted 10 times in 100% ethanol and sonicated again. 20 μl of the cells were placed on stubs using glass coverslip and kept for drying in a CO_2_ desiccator for overnight. After the overnight drying, the stubs containing cells were gold coated using sputter coater and visualized in HR FESEM (High-resolution field emission scanning electron microscope) with a resolution of 1.0 nm at 15 kV.

### 2.7. Immunoblotting Analysis

The 5 ml cultures of WT and different mutants of yeast were grown at 24°C until 1.5 OD_600_, harvested by centrifugation at low speed, washed once with 20% TCA and stored at −80°C freezer until they were used. The whole cell protein lysates were prepared from the frozen cells by 20% Trichloroacetic acid (TCA), and immunoblotting was conducted. In brief, cell pellets were re-suspended in 200 μl of 20% TCA and lysed in the presence of glass beads by vortexing for 20 min at room temperature, centrifuged for 10 min at 7K rpm, and pellets were collected and washed once with 1.0 ml 0.5M Tris-HCl, pH7.5. The washed pellets were re-suspended in 200 μl of 0.5M Tris-HCl, pH7.5; SDS loading dye was added, boiled for 5 min and analyzed by SDS-PAGE. Proteins were transferred onto Nitrocellulose membrane for western blotting. Blots were probed with following primary antibodies were used: Anti-Slt2-P/phospho-p42/44 MAPK (catalog No. 4370; Cell Signaling), Anti-Mpk1 (yC-20-catalog No. sc-6803; Santa Cruz), and Anti-Myc (9E10). The IR-Dye 800CW Anti-Rabbit IgG, Anti-Goat, and Anti-Mouse were used as secondary antibodies for respective primary antibodies. Polyclonal antibodies against recombinant TBP was raised in rabbit. Western blots were scanned by using an Odyssey infrared imager (LI-COR Biosciences).

### 2.8. Observation and analysis of flocculation

Overnight grown yeast cells were re-suspended to an equal cell density in SC media. Flocculation was observed using a method mentioned earlier (Church *et al.* 2017) with some modifications. In brief, 3 ml of cell cultures of 2 OD_600_ were placed into a tissue culture plate. Cells were subsequently agitated by shaking and then left undisturbed. Images of plates were subsequently recorded by HP scanner at time 0, 15 and 30 minutes. The % Flocculation was measured by using a method described earlier (Bony *et al.* 1998). Briefly, yeast cells were deflocculated by two washes with wash buffer (50mM sodium acetate, pH 4.5 and 5mM EDTA) and twice with distilled water. Cells were re-suspended in flocculation buffer (50mM sodium acetate, pH 4.5 and 5mM calcium chloride) and incubated for 60 min at 100 rpm in the incubator shaker at RT and then 6 mL of cell suspension was taken into culture tubes and kept vertically undisturbed for 10 min to allow cells for settling. The following equation determined percentage flocculation ability (F): F= (A-B/A) × 100% as described previously (Singh *et al.* 2015). ‘A’ indicates initial OD before shaking at time=0, and ‘B’ indicates final OD after 70 min of incubation (60 min in shaking + 10 min settling time). The percentage of flocculation is represented as mean of two independent biological replicates.

### 2.9. Isolation of total RNA and RT-qPCR

Total RNAs were extracted using a standard method (Schmitt *et al.* 1990). The cDNA preparations were performed by using an iScript cDNA synthesis kit procured from BioRad. For PCR amplification, 2X SYBR green master mix was used. The conditions for PCR amplification were as follows: initial denaturation for 3 min at 95°C, PCR amplification for 40 cycles with denaturation for the 20s at 95°C, annealing for 20s at 58°C and elongation at 72°C for 20s in an ABI 7300 Real-Time PCR machine (Applied Biosystems). The relative expression levels (mRNAs) of each gene in wild-type and mutants were normalized by subtracting the β-actin threshold cycle (CT) values and the fold change (increase or decrease) was calculated through the 2^-ΔΔCT^ method (Livak and Schmittgen 2001). Three independent experiments were performed, and each sample was run in triplicate. Primers used in this study are listed in Supplementary Table S2.

### 2.10. Chromatin Immunoprecipitation

The ChIP assays were performed as described earlier (Tomar *et al.* 2008). Cells were grown in 100 ml of SC growth media until 1.2 to 1.4 OD_600_. Subsequently formaldehyde was added (1% final concentration) for crosslinking by incubation for 10 min, and then 6ml of 2.5M glycine was added to stop the crosslinking reaction by incubation for 5 min at 25°C, cells were harvested by centrifugation, washed by water and stored in −80°C freezer until they were used. For preparation of soluble chromatin extracts, frozen cell pellets were lysed by vortexing for 30 min at 4°C in presence of glass beads in 1.2 ml of FA lysis buffer (50 mM HEPES, pH7.5, 150 mM NaCl, 2mM EDTA, pH8.0, 1% Triton X-100, 0.1% Sodium Deoxycholate and 0.1% SDS), glass beads were removed by centrifugation and final volume was made up to 1.8 ml by adding FA lysis buffer. The lysed cells were then sonicated to shear the chromatin into fragments averaging 200 - 400 bp using a Bioruptor (Diagenode) sonicator. The sonication conditions were as follows; high power, 8 cycles with the 30s ON and OFF in ice-chilled cold water. The soluble chromatin of sonicated cell lysates was collected by centrifugation at 14K RPM for 30 min at 4°C. For ChIP assay, 100 μl of chromatin extracts were incubated with 1μl of Anti-TBP, Anti-Tup1, Anti-Pol-II (ab817, 8WG16) and Anti-Myc (Rlm1) antibodies for overnight. The Anti-TBP and Anti-Tup1 antibodies were generous gifts from Dr. Joseph Reese. Multiple dilutions of input DNA were used to find out the amount of template DNA for PCR amplification to be in linear range. The immune-precipitated DNAs were amplified by RT-qPCR using SYBR Green master mix and ABI 7300 PCR machine. The following primer sets, specific for upstream regions of *FLO1*; *FLO1P1* (−215/+14), *FLO1P2* (−431/-312), *FLO1P3* (−612/-457) and *FLO1P4* (−817/-659) and *FLO5*; *FLO5P1* (−139/+20), *FLO5P2* (−413/-264), *FLO5P3* (−621/-434) and *FLO5P4* (−804/-595) were used. The *STE6* amplicons were used for normalization, as an internal control. The relative fold change was calculated by using the 2^-ΔΔCT^ protocol. The results presented here are the averages of three independent biological repeats.

### 2.11. Statistical analysis

Statistical analyses of the data were performed using GraphPad Prism 5.0 software of three independent repeats. Results of statistical analyses were presented as means ± standard deviations (SD). Differences between groups were tested using Student’s *t*-test, with *P* values of <0.05 were considered as statistically significant. Asterisks used in the figures represent the following significance values: \**P* ≤ 0.05; \*\**P* ≤ 0.01; \*\*\**P* ≤ 0.001.

## 3. Results

### 3.1. Mutations in *SEN1* induces enlarged cell size, rough surface morphology, and abnormal cell wall chitin content

We have earlier shown de-repression of *FLO* genes in mutants of *Sen1, Nab3* and *Rnt1* cells of yeast which leads to calcium-dependent flocculation phenotype (Singh *et al.* 2015). Recently one study reported the upregulation of *FLO1* in a *Sen1* mutant strain suggesting the contribution of Sen1 in the regulation of flocculation phenotype (Chen *et al.* 2017). Another study has shown a correlation between higher cell wall chitin content and sensitivity to cell wall damaging agents (Liu *et al.* 2010). However, a molecular mechanism connecting the CWI pathway to flocculation needs to be elucidated. The cell wall of yeast remains in direct contact with the surrounding environment and regulates; cell size, cell shape, cell-cell, and cell-surface interactions. Many of the studies have suggested that cell-cell interactions are majorly facilitated by cell wall adhesin proteins, also known as flocculins which are responsible for flocculation phenotype (Barua *et al.* 2016; Bester *et al.* 2012). To elucidate the role of Sen1 in the mechanism of flocculation, we performed a series of experiments to find whether or not flocculation phenotype is connected with cell wall composition, cellular morphology and growth response in the presence of cell wall damaging agents. First; we performed microscopy and SEM experiments for morphological analysis. Second; the FACS experiment was conducted to measure cell wall chitin content using fluorescent CFW stain and cell size by plotting values of forward versus side scattering of wild-type, non-flocculating and flocculating strains. Third; the effect of cell wall damaging agents (CongoRed, CFW and caffeine) on the growth of cells was analyzed by spot assay and counting CFUs. Our careful analysis of microscopy and FACS results suggested that the flocculating mutant cells as compared to wild-type possess enlarged cell size and distinct rough surface morphology (Figure 1A and B) indicating defect in the cell wall. Strikingly we observed a higher level of chitin in the flocculating *Sen1* mutant strains as compared to wild-type and non-flocculating mutant cells (Figure 1C). In spot assays, we found that most of the flocculating *Sen1* mutants (*sen1-1*, *sen1 ∆N*, and *sen1-K128E*) which display abnormal cell size and surface morphology, showed growth defect, suggesting that Sen1 plays an essential role in cell wall maintenance (Figure S1 and S2). The early studies revealed that Sen1 is one of the main components of a protein complex known as NNS (Nrd1, Nab3, and Sen1), possess RNA/DNA helicase activity (Martin-Tumasz and Brow 2015) and functionally linked with factors of RNA processing machinery; Rrp6, TRAMP/Exosome and Rnt1 (Fasken *et al.* 2015; Finkel *et al.* 2010; Fox *et al.* 2015). We therefore, extended our investigations with available mutants of Nrd1 (*nrd1-102*), Nab3 (*nab3-11*), deletion mutants of TRAMP (*air1∆, air2∆, trf4∆*)*, rrp6∆* and *rnt1∆* cells. Strikingly we detected higher chitin content in *rnt1∆* and *nab3-11* mutants as well. However, the cell size and chitin content of TRAMP and Nrd1 mutants was found similar to wild-type cells indicating that the role of Sen1 in regulation cell wall maintenance is not dependent TRAMP and Nrd1 (Figure S3A and S3B). Furthermore, as Sen1 has been shown to interact with Rnt1 (Finkel *et al.* 2010; Ursic *et al.* 2004) and Nab3 as part of NNS complex (Vasiljeva *et al.* 2008), we believe that role of Sen1 in cell wall integrity is probably dependent on Nab3 and Rnt1. Our comparative analysis of flocculating and non-flocculating cells indicate that only strongly flocculating mutant strains showed enlarged cell size, rough surface morphology, and abnormal chitin content. These studies demonstrate that yeast Sen1 in association with Nab3 and Rnt1 plays an essential role in the regulation of flocculation phenotype by cell wall maintenance.

**Figure 1:**
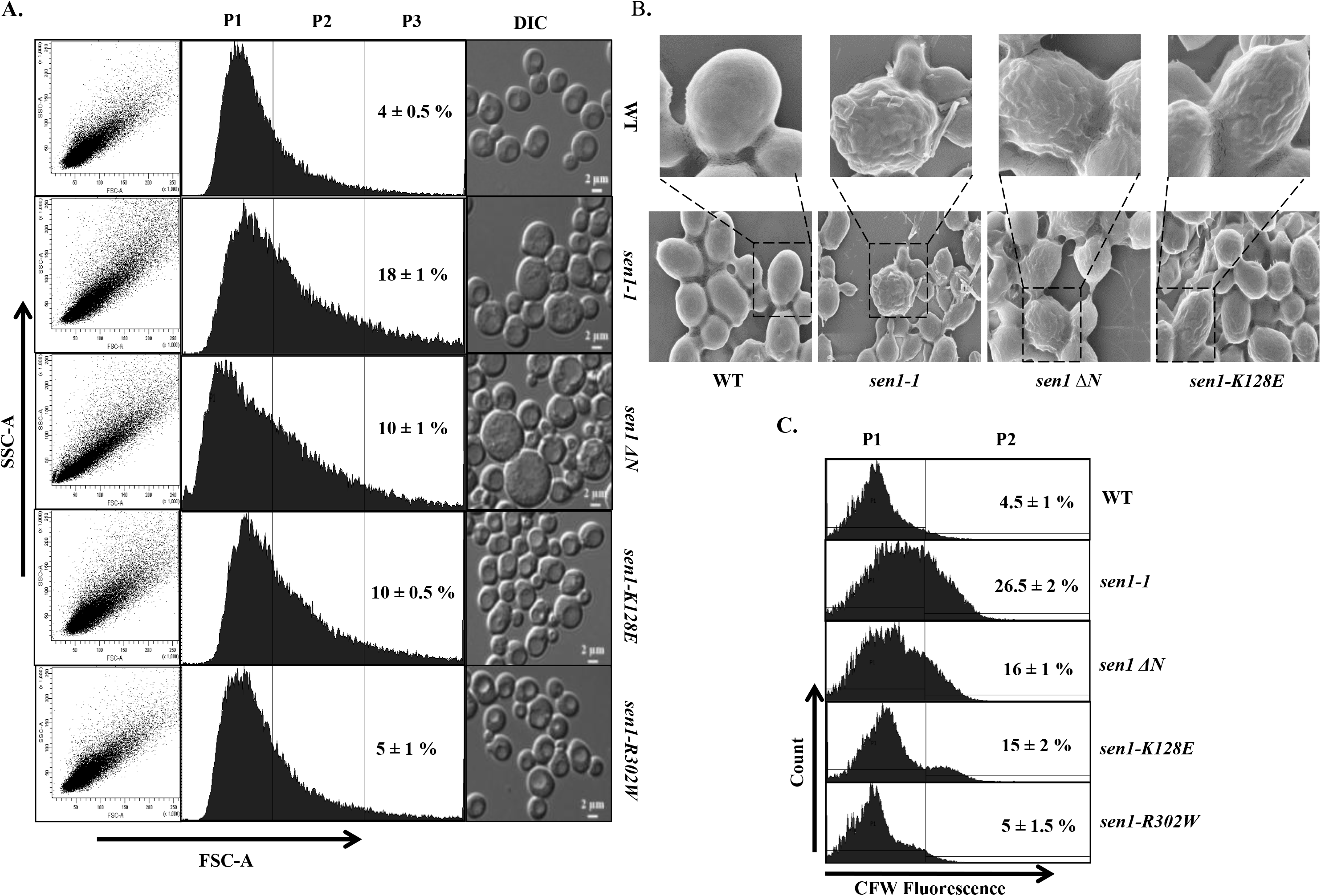
Cell wall defect correlates with abnormal cell size, cellular morphology, and chitin content. A. Cell size measurement of wild-type and *Sen1* mutants (*sen1-1, sen1 ΔN, sen1-K128E* and *sen1-R302W*) by plotting values of SSC vs. FSC obtained from FACS analysis by using BD-FACS Area III instrument. Cells at 0.8 OD_600_ were harvested, washed and re-suspended for FACS analysis. Equal numbers of cells (30000 events) were gated, and scattering was recorded. Plots of forward and side scattering were obtained and analyzed. P3 gate indicates the percentage of large cells. Representative DIC images of cells as indicated in figure acquired by using a Carl Zeiss motorized Apotome fluorescence microscope showing the size of each cell (scale bar 2 μm) are presented on the right side of respective plots.
B. The saturated cultures of cells (WT, *sen1-1, sen1 ∆N* and *sen1-K128E*) were harvested, washed, fixed with 4% Paraformaldehyde for 5 min, washed twice with PBS and dehydrated sequentially with increasing concentration of ethanol (30-100%). Cells were subsequently pulse sonicated and diluted 10 times in 100% ethanol. 20 μl of diluted cells were placed on stubs for O/N drying in a CO_2_ desiccator; gold coating was done by sputter coater and images were captured by using Zeiss High-Resolution Field Emission Scanning Electronic Microscope (HR-FESEM).
C. An equal number of exponentially growing cells (1 ml culture) as indicated in the figure were stained with CFW (50μg/ml) for 30 min, harvested, washed, re-suspended in PBS and analyzed by FACS. Equal numbers of cells (30000 events) were used for fluorescence using DAPI filter. P2 gate indicates the percentage of cells having increased cell wall chitin.

### 3.2. Mutations in *SEN1*, *NAB3*, and deletion of *RNT1* induces constitutive activation of CWI pathway

In budding yeast, the Cell Wall Integrity (CWI) signaling pathway which composed of an evolutionarily conserved kinase cascade is majorly responsible for maintenance of cell wall homeostasis (Jimenez-Sanchez *et al.* 2007; Levin 2005; Levin 2011). The CWI pathway regulates the transcription of enzymes, required for cell wall synthesis under a variety of stress conditions including unfolded protein response (Scrimale *et al.* 2009) and lipid homeostasis. Many studies have shown that during cell wall stress, different cell membrane sensor proteins interacts with the guanine nucleotide exchange factor (Rom2) leading to the activation of small GTPase (Rho1) which activates protein kinase C (Pkc1) as shown in the schematic (Figure 2A). Pkc1 transmits the signal to a MAPK module, MAPKKK (Bck1), MAPKK (Mkk1 and Mkk2), and the MAPK (Slt2/Mpk1). The activated Slt2 is known to trigger the cell wall damage-specific transcriptional response by activating Rlm1, a transcription factor (Bermejo *et al.* 2008; Sanz *et al.* 2012). The Mpk1/Slt2 has also been shown to prevent the recruitment of Sen1-Nrd1-Nab3 termination complex to regulate the expression of CWI pathway genes (Kim and Levin 2011). The cell wall chitin through covalent binding with glucan plays an essential role in controlling the morphogenesis and fungal cell wall remodeling which is necessary to counterbalance the cell wall stress (Arroyo *et al.* 2016; Ene *et al.* 2015). Since flocculating mutant strains showed higher chitin content, enlarged cell size and sensitivity to cell wall damaging agents, we first decided to measure the phosphorylation of Slt2 by quantitative western blotting using an antibody specific to the phosphorylated form of Slt2. Interestingly we found constitutive phosphorylation of Slt2 (8-10 fold increase) in all flocculating *Sen1* mutant strains*; sen1-1*, *sen1 ∆N*, *sen1-K128E* (Figure 2B and 2C). However, the phosphorylation of Slt2 was also induced in *nab3-11, rnt1∆* and *nrd1-102* but lesser than flocculating *Sen1* mutants in comparison to wild-type and a non-flocculating *Sen1* mutant (*sen1-R302W*), TRAMP mutants (*air1∆, air2∆, trf4∆*) and *rrp6∆* (Figure S4A and S4B). Western blot experiment to measure Slt2 phosphorylation was repeated 3-4 times, only one of the representative images is presented here. Western blot results of two biological repeats to measure Slt2 phosphorylation in *Sen1* mutant strains are available in the supplementary file (Figure S4C). Previously, we reported the flocculation phenotype of few of the mutants of *Sen1, Nrd1, Nab3*, and *rnt1∆*, but the connection with CWI pathway was not established. These results suggest that the activation of the CWI pathway is functionally linked with flocculation of yeast cells.

**Figure 2:**
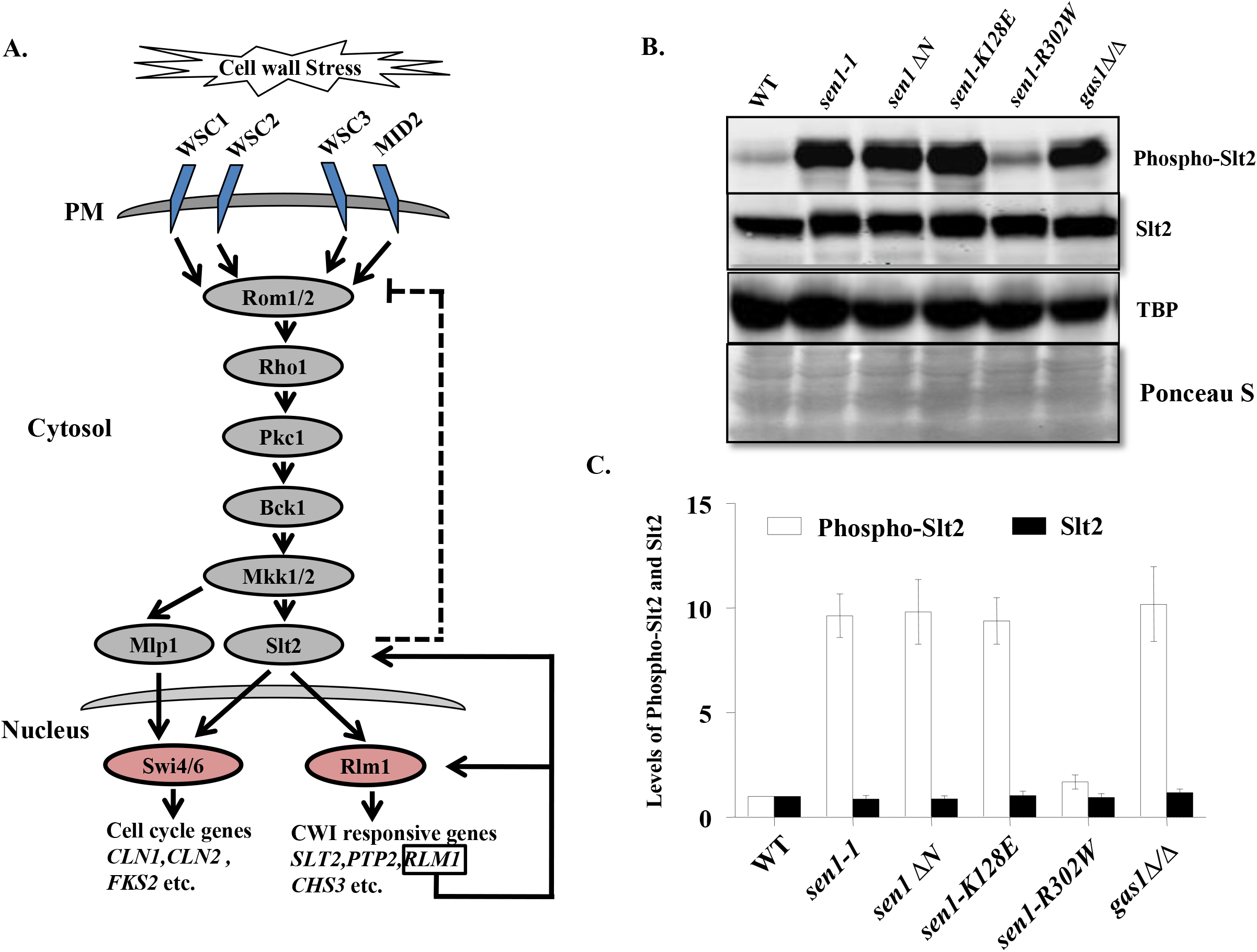
Flocculating *Sen1* mutants shows constitutive phosphorylation of Slt2. A. Schematic representation of the Cell Wall Integrity (CWI) pathway. Under cell wall stress conditions, sensor proteins transmit the signal to Rom1/2 G-protein which in turn activates Pkc1 leading to the activation of MAP Kinase cascade consisting of Bck1, Mkk1/2, and Slt2. Activated Slt2 enters into the nucleus to activate Rlm1, a transcription factor which binds to responsive genes to induce the expression.
B. Exponentially growing cultures of wild-type, flocculating (*sen1-1, sen1 ∆N* and *sen1-K128E*) and non-flocculating (*sen1-R302W*) mutants of *Sen1* were harvested, washed and whole-cell extracts were prepared by 20% TCA as described in material and methods. Cell extracts were separated on 10% SDS-PAGE and transferred onto nitrocellulose membrane. Blots were probed with primary antibodies as indicated for checking levels of phosphorylated and non-phosphorylated forms of Slt2. For protein loading controls, blots were re-probed with anti-TBP antibody and stained with Ponceau S as indicated.
C. Quantification of phosphorylated and non-phosphorylated forms of Slt2 relative to the wild-type and normalized by TBP using ImageJ software. Bars in this data represent the difference in Slt2 phosphorylation in each mutant compared to that of wild-type cells. Error bars represent SD. Data shown here are the averages of three independent experiments.

### 3.3. The phosphorylation of Slt2 correlates with de-repression of *FLO* genes and flocculation phenotype

It is now clear that flocculating *Sen1* mutant cells (*sen1-1*, *sen1 ∆N*, and *sen1-K128E*) exhibit constitutive increase in phosphorylation of Slt2 and alteration in cell wall integrity pathway as they are sensitive to cell wall damaging agents (Figure S1A) and enlarged cell size. Thus we decided to understand the underlying mechanism further. Under cell wall stress conditions, Slt2 is phosphorylated to induce the expression of responsive genes. Therefore, first, we measured the Slt2 phosphorylation in wild-type cells upon cell wall damage. We treated the cells with cell wall damaging agent (CFW) and exposed to a higher temperature to artificially induce the cell wall damage. The growth of wild-type cells was not affected at a higher temperature as well as in the presence of CFW (Figure S5A, S5B and Figure 3A, 3B), whereas, the phosphorylation of Slt2 was significantly induced in both these conditions (Figure 3C). Next, to investigate whether phosphorylation of Slt2 has any correlation with the expression of *FLO* genes, we performed the RT-qPCR to measure the mRNA expression of *FLO1, 5* and *9.* Interestingly we observed 3-5 fold increase in expression of *FLO1, FLO5* and *FLO9* (Figure 3D) than untreated cells, leading to the induction of flocculation/aggregation of cells upon stress induced by temperature and CFW treatments (Figure 3E). To further understand the role of CWI pathway in expression of *FLO*s (*1, 5, 9* and *10*), we measured the basal level mRNA expression by RT-qPCR of these genes in deletion mutant of CWI pathway; *wsc1∆/∆, wsc2∆/∆, wsc3∆/∆, mid2∆/∆, rom2∆/∆, bck1∆/∆, mlp1∆/∆, slt2∆/∆, rlm1∆/∆* and double deletion mutant, *slt2∆/∆ mlp1∆/∆*. We detected about 2-3 fold decrease in basal expression of *FLO* genes in almost all the mutants except *mlp1∆/∆* cells which further suggest that CWI signaling pathway regulates the expression of *FLO* genes (Figure 3F). The double deletion mutant (*slt2∆/∆ mlp1∆/∆*) was utilized to test whether the signal is transmitted through Mlp1 or Slt2. Interestingly we noticed a decrease in transcription of *FLO* genes in double deletion mutant than single deletion (*mlp1∆/∆* indicating that transcription is regulated through Slt2-Rlm1, not via Mlp1-Swi4/6 branch. A schematic for the entire CWI signaling pathway has been shown in Figure 2A. Thus our results suggest that the activation of CWI pathway is essential for eliciting the expression of *FLO* genes and flocculation phenotype which provides a suitable microenvironment required for cell survival under stress conditions (Goossens *et al.* 2015).

**Figure 3:**
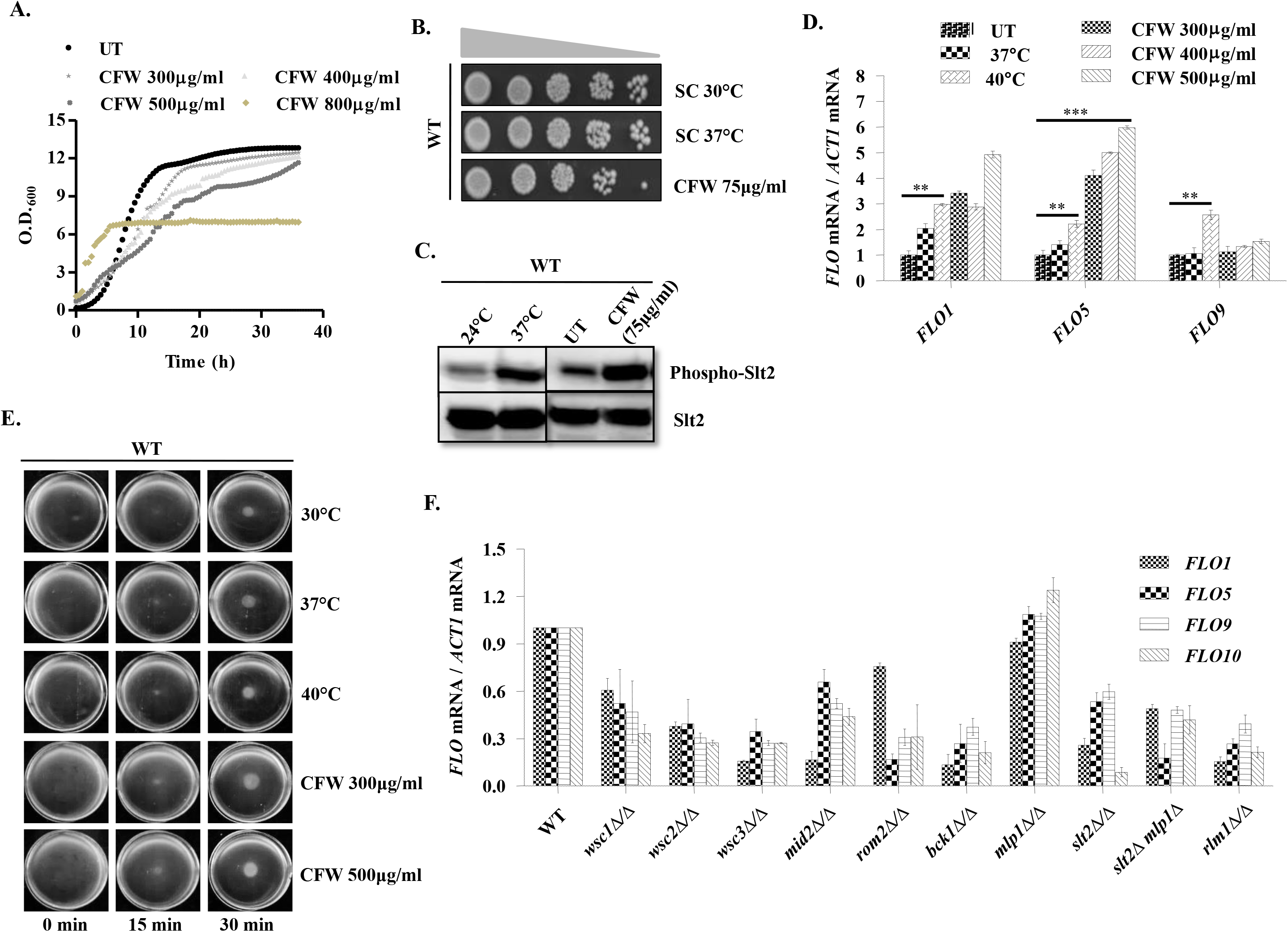
Cell Wall Integrity pathway is required for the expression of *FLO* genes. A. Growth curve analysis of wild-type cells was measured in liquid growth media in absence and presence of CFW concentrations as indicated in an automatic spectrophotometer plate reader by taking OD_600_ at regular intervals of 30 min for 36 hours.
B. Spot assay was performed to test the growth of wild-type cells in the presence of CFW and at 37°C. Cells were grown O/N, diluted to 1 OD_600_ in 1 ml of water and diluted further 4 times (10 fold serial dilution). 3 μl of cells from each dilution were spotted on SC-Agar plates with and without CFW, incubated at 30°C. For temperature stress, cells were spotted on SC-Agar plates and incubated at 37°C. Growth was recorded by scanning after 72 hours of incubation.
C. Wild-type cells were grown until 0.8 OD_600_ and grown further for 3 hours in the presence of CFW as well as at 37°C as indicated to induce cell wall stress. Whole cell extracts were prepared by 20% TCA and phosphorylation of Slt2 were measured by quantitative western blotting.
D. Quantification of mRNAs of *FLO1*, *5*, and *9* by RT-qPCR. The 5 ml cultures of exponentially growing wild-type cells were grown for 3 hours at 30°C, 37°C, 40°C and in the presence of CFW concentrations as indicated at 30°C for 3 hours. The cells were subsequently harvested, total RNAs were extracted by ‘heat phenol freeze method’ as described in materials and methods and cDNA was prepared by using an iScript cDNA synthesis kit purchased from Biorad. The gene-specific primers were used for amplification of mRNAs by RT-qPCR. *ACT1* mRNAs were used as an internal control for quantification of *FLO* mRNAs.
E. Images showing flocculation of wild-type yeast cells grown O/N at 30°C, 37°C, 40°C and in the presence of CFW. O/N grown cultures were diluted to 2 OD_600_ in 3 ml volume of fresh SC growth media and poured into 35 mm Petri dishes. The cells in plates were agitated for uniform distribution by shaking and then left uninterrupted; subsequently, pictures were recorded at time 0, 15 and 30 min.
F. Quantification of mRNAs of *FLO1, 5, 9* and *10* by RT-qPCR. Exponentially growing cultures at 30°C of wild-type and deletion mutant cells of the CWI pathway as indicated were harvested, total RNAs were extracted, and cDNA was prepared for RT-qPCR to measure the expression of *FLO* genes. *ACT1* mRNAs were used as internal control for quantification. Bars in this data represent the difference in fold change of *FLO* gene expression in WT cells compared to that of untreated and treated cells. Error bars represent SD. Data shown here are the averages of three independent experiments. Statistical analysis was carried out with a two-tailed, unpaired, Student’s *t*-test to analyze differences between the untreated to the Temp at 40°C and CFW treated (500μg/ml) of wild-type cells:\*\**P* ≤ 0.01; \*\*\**P* ≤ 0.001.

### 3.4. Deletion of *SLT2* suppresses the expression of *FLO* genes and flocculation

We have so far observed a strong correlation between Slt2 phosphorylation and expression of *FLO* genes. To further validate this observation, we deleted the *SLT2* from two of the flocculating *Sen1* strains (*sen1-1* and *sen1-K128E*) to create double mutants which were confirmed by western blotting using an antibody specific to Slt2 protein (Figure 4A). The wild-type and mutant strains (*sen1-1* and *sen1-K128E*) with and without *SLT2* were utilized to measure the mRNA expression by RT-qPCR of *FLO1, 5, 9* and *10* genes. As expected we observed a drastic decrease in expression of *FLO* genes in the absence of *SLT2* measured by RT-qPCR (Figure 4B) leading to the suppression of flocculation phenotype, examined by recording cell aggregation in Petri dishes at indicated time points (Figure 4C). Furthermore, flocculation of cells was quantified as % flocculation which was also decreased in the absence of *SLT2* in *sen1-1* and *sen1-K128E* (Figure 4D). Moreover, the growth of the wild-type and *Sen1* mutant strains with and without *SLT2* was tested by spot assay in the presence of cell wall damaging agents; CongoRed, CFW and Caffeine. Interestingly, *sen1-1 slt2∆* and *sen1-K128E slt2∆* cells were found more sensitive to the cell wall damaging agents in comparison to wild-type, *sen1-1* and *sen1-K128E* cells (Figure S6). However, the *slt2∆* cells were found more sensitive than wild-type, *sen1-1* and *sen1-K128E* but less than *sen1-1 slt2∆* in the presence of cell wall damaging agents. These observations suggest that Slt2 dependent regulation of *FLO* genes is required for survival under cell wall stress conditions.

**Figure 4:**
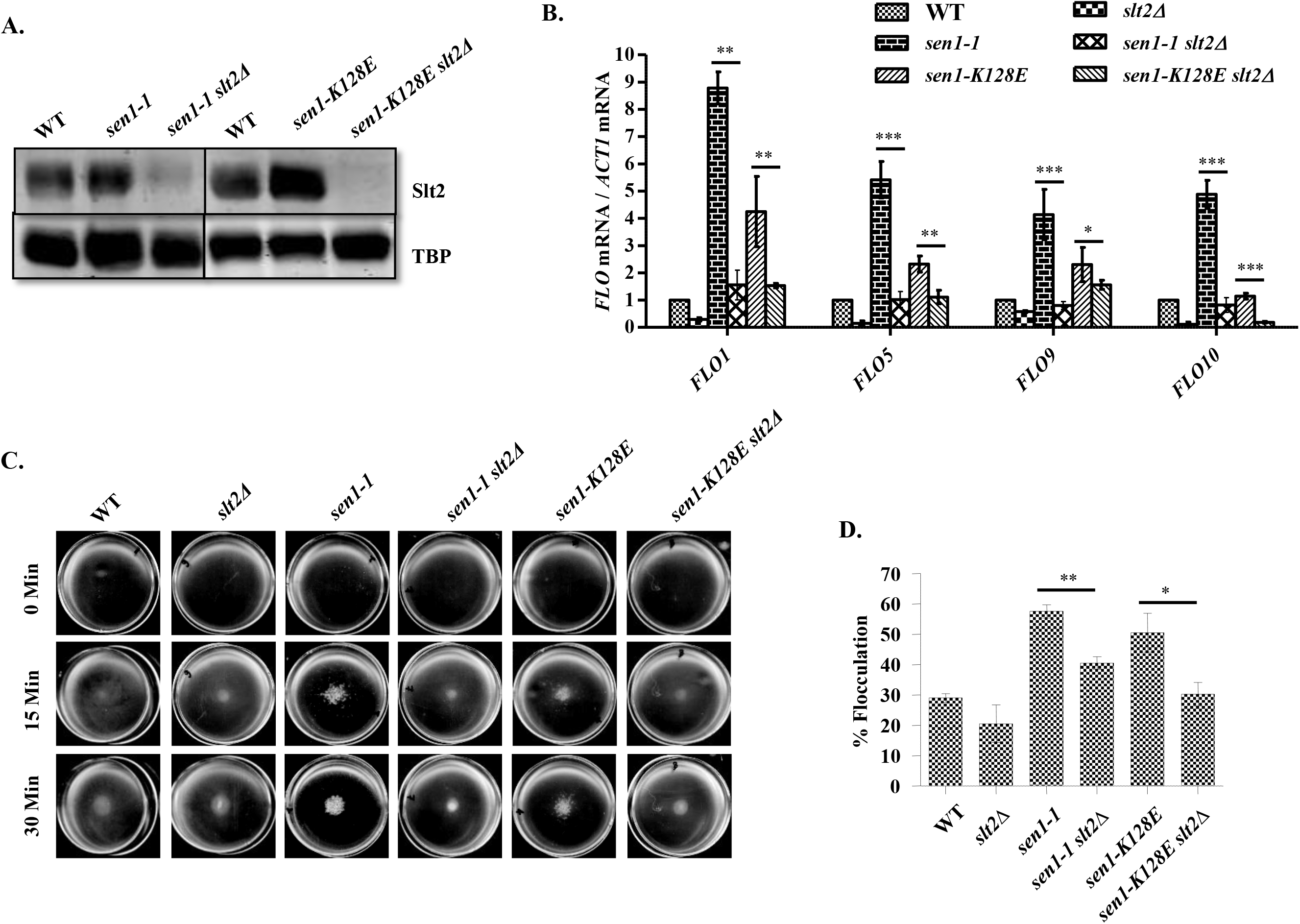
Slt2 is required for expression of *FLO* genes. A. Confirmation of *SLT2* deletion in WT, *sen1-1* and *sen1-K128E* cells by western blotting. TBP western was performed for loading control.
B. The mRNA quantification of *FLO1, 5, 9* and *10* by RT-qPCR. Cells were grown at 30°C, total RNAs, and cDNAs were prepared. *ACT1* mRNAs were used as internal control for quantification of *FLO* mRNAs.
C. Flocculation of wild-type and *Sen1* mutant cells (*sen1-1* and *sen1-K128E*) with and without *SLT2* was recorded by taking pictures at indicated time points as described earlier.
D. The % flocculation of wild-type and *Sen1* mutant cells (*sen1-1* and *sen1-K128E*) with and without *SLT2* was calculated. Bars in this data represent the difference in fold change of *FLO* gene expression (C) and % flocculation (D) in each mutant compared to that of wild-type cells. Error bars represent SD. Data shown here are the averages of three independent experiments. Statistical analysis was carried out with a two-tailed, unpaired, Student’s *t*-test to analyze differences between the *sen1-1* and *sen1-K128E* with and without *SLT2* deletion:\**P* ≤ 0.05; \*\**P* ≤ 0.01; \*\*\**P* ≤ 0.001.

### 3.5. The binding of Rlm1 correlates with upregulation of *FLO* genes and flocculation phenotype

The activation of the CWI pathway occurs in cell wall damage conditions. A few years ago the role of Slt2-dependent CWI pathway was suggested in cell wall remodeling and filamentous growth of fungus (Birkaya *et al.* 2009) which further supports our observations. Phosphorylation of Slt2 is the key event in the activation of the CWI signaling pathway leading to the recruitment of Rlm1 and SWI2/SNF2 to regulate the expression of CWI responsive genes (Sanz *et al.* 2012). As our results reveal constitutive phosphorylation of Slt2 in flocculating strains, we hypothesized that it might lead to the recruitment of Rlm1 at the promoters of *FLO* genes. First, we searched for the Rlm1 binding sites at the *FLO* genes through sequence analysis. A consensus nucleotide sequence, **TA(A/T)_4_TAG** was identified as the Rlm1 binding sequence by site selection experiments using random oligonucleotide sequences (Dodou and Treisman 1997; Jung and Levin 1999). Interestingly our sequence analysis revealed similar recognition sequences at the upstream regions of *FLO1, FLO5, FLO9* and *FLO10* promoters (Figure 5A) suggesting that Rlm1 probably binds at the promoters of *FLO* genes to activate the transcription. To investigate whether Rlm1 is recruited to the *FLO* promoters or not, we performed chromatin IP experiments. Chromatin extracts were prepared from formaldehyde crosslinked Rlm1 Myc tagged strains; wild-type, *sen1-1* (flocculating strain) and a non-flocculating mutant, *sen1-R302W*. The Anti-Myc antibody was used for the immunoprecipitation. Four primer sets for *FLO1* upstream sequences (ranging from +14 to −817) and *FLO5* (ranging from +20 to −804) were used for PCR amplification of Rlm1 ChIP DNA (Figure 5B). Our analysis also suggests that Rlm1 binding sites overlap with the Tup1 binding site (Fleming *et al.* 2014) indicating that these two proteins probably compete for binding to regulate the expression of *FLO* genes. We found around 2.5 fold enrichment of Rlm1 at the *FLO1*, and *FLO5* analyzed by RT-qPCR using Rlm1 ChIP DNA and primer sets for *FLO1* and *FLO5* as indicated in the schematic. The occupancy of Rlm1 was relatively found more in *sen1-1* flocculating cells than wild-type and non-flocculating *Sen1* mutant, suggesting that Rlm1 is constitutively recruited at the *FLO* genes (Figure 5C). This result provides strong support to our hypothesis that Rlm1 physically binds at the promoters of *FLO* genes to regulate their expression. We observed peak PCR amplification of Rlm1 chip DNA with P2 and P3 primer sets, suggesting that recruitment of Rlm1 majorly occurs at around −300 to - 600 bp of *FLO1* and *FLO5*. Primers for the *STE6* gene were used as internal control for the calculation of fold change. Further to study the role of Rlm1 in the expression of *FLO* genes and flocculation phenotype, we first deleted the *RLM1* gene from wild-type, flocculating *Sen1* mutant strains (*sen1-1*, *sen1 ∆N*, and *sen1-K128E*) and non-flocculating *Sen1* mutant, *sen1-R302W*. Interestingly upon deletion of *RLM1*, we detected a noticeable decrease in flocculation/aggregation of cells measured by taking pictures at indicated time points of cell suspensions placed in Petri-dishes (Figure 5D). The % flocculation calculated by taking OD_600_ at different time points during the settling of cells in glass tubes was also reduced after deletion of *RLM1* (Figure 5E). Subsequently, RT-qPCR was performed to quantify the expression of *FLO* genes in all these strains in the presence and absence of *RLM1*. As expected, we found good reduction (~50%) in the expression of mRNAs of *FLO* genes in *sen1-1*, *sen1 ∆N*, and *sen1-K128E* strains after deletion of *RLM1* indicating that Rlm1 acts as a transcription factor for the expression of *FLO* genes (Figure 5F). Furthermore, the effect of cell wall damaging agents on the growth of cells after *RLM1* deletion as well as phosphorylation of Slt2 was examined. To this end the inhibitory effect of cell wall damaging agents on the growth was suppressed after deletion of *RLM1* in contrast to deletion of *SLT2*, suggesting that CWI pathway dependent expression of *FLO* genes is essential for survival under stress conditions (Figure S7A). Since Slt2 is located at the upstream of Rlm1 in the signaling pathway, we did not detect any significant reduction in phosphorylation of Slt2 after *RLM1* deletion (Figure S7B and S7C). A slight decrease in Slt2 protein level was observed because Rlm1 also acts as the transcription factor for expression of Slt2 through a regulatory feedback mechanism (Garcia *et al.* 2016). A molecular mechanism for the recruitment of Rlm1 and regulation of *FLO* genes has been investigated further as described from now onward in this manuscript.

**Figure 5:**
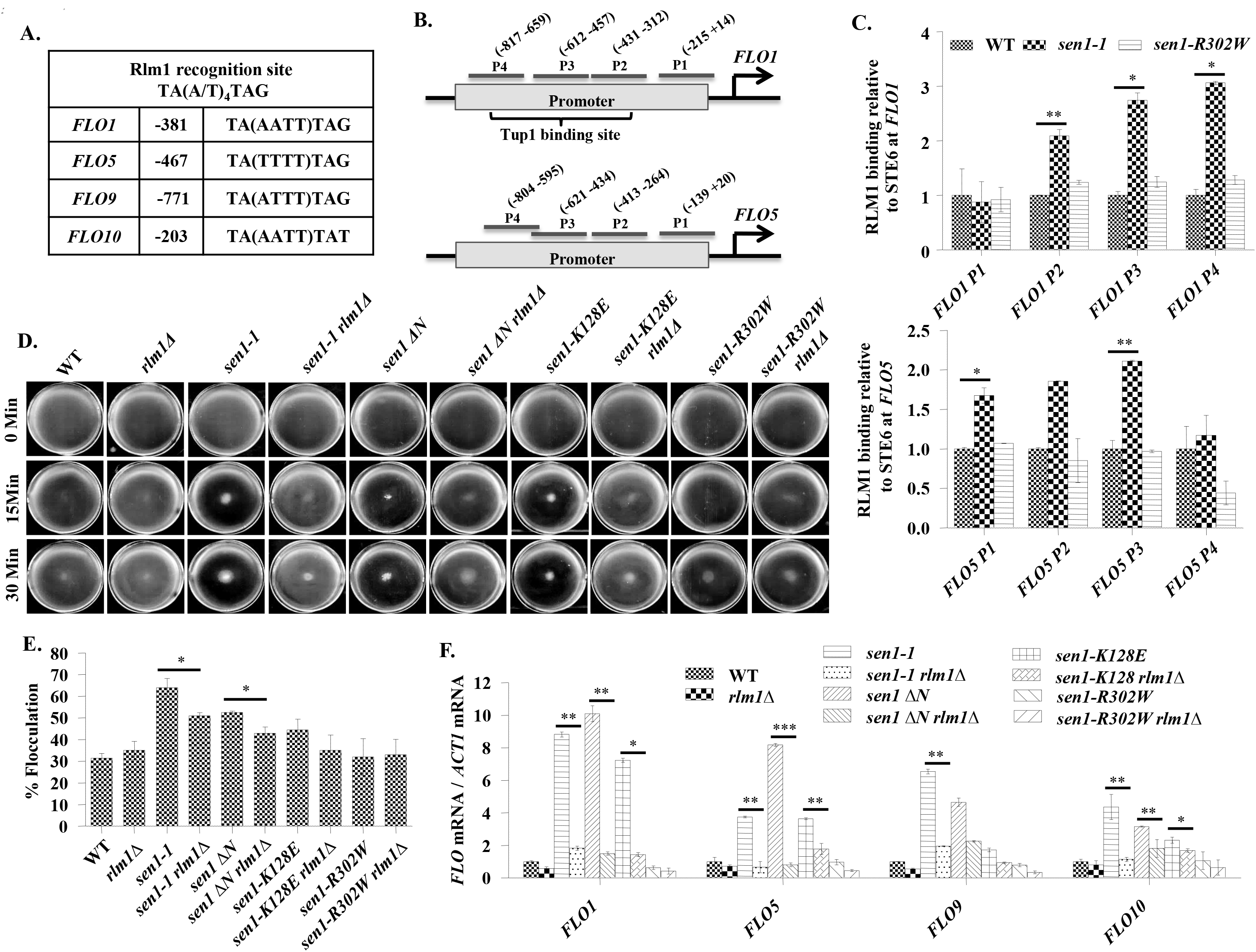
Rlm1 binding at the *FLO* genes is required for their expression. A. Identification of Rlm1 binding sites at the upstream regions of *FLO* genes (*FLO1, 5, 9* and *10*).
B. Schematic representation of *FLO1* and *FLO5* promoters showing primer sets designed for upstream regions including Rlm1 binding site used for Chromatin Immunoprecipitation (ChIP) analysis. Tup1 binding is also indicated in the schematic, adapted from (Fleming *et al.* 2014).
C. ChIP assay was performed as described in materials and methods with chromatin extracts isolated from Rlm1 Myc tagged wild-type, *sen1-1*, and *sen1-R302W* cells. Anti-Myc antibody was used for immunoprecipitation. ChIP DNAs were analyzed by RT-qPCR. Primers of *STE6* gene were used as internal control for quantification of Rlm1 occupancy at P1, P2, P3 and P4 upstream locations of *FLO1* and *FLO5* genes.
D. Flocculation of wild-type and *Sen1* mutant cells (*sen1-1, sen1 ∆N, sen1-K128E* and *sen1-R302W*) with and without *RLM1* was recorded by taking pictures at indicated time points.
E. The % flocculation of wild-type and *Sen1* mutant cells (*sen1-1, sen1 ∆N, sen1-K128E*, and *sen1-R302W* with and without *RLM1* were calculated.
F. The mRNA quantification by RT-qPCR of *FLO1, 5, 9* and *10* genes in wild-type and *Sen1* mutant cells (*sen1-1, sen1 ∆N, sen1-K128E* and *sen1-R302W*) with and without *RLM1*. Cells were grown at 30°C, total RNAs, and cDNAs were prepared. *ACT1* mRNAs were used as internal control for quantification of *FLO* mRNAs. Bars in this data represent the difference in fold change of Rlm1 binding (C), % flocculation (E) and *FLO* gene expression (F) in each mutant compared to that of wild-type cells. Error bars represent SD. Data shown here are the averages of three independent experiments. Statistical analysis was carried out with a two-tailed, unpaired, Student’s *t*-test to analyze differences between the indicated mutants and the wild-type strain (C) and between mutants with and without *RLM1* deletion (E and F): \**P* ≤ 0.05; \*\**P* ≤ 0.01; \*\*\**P* ≤ 0.001.

### 3.6. Binding of Rlm1 reduces the occupancy of Tup1 leading to the recruitment of TBP and Pol-II at the promoters of *FLO* genes

Previous studies have shown that reduced occupancy/deletion of *TUP1* results in strong de-repression of *FLO* genes (Fleming *et al.* 2014). It was also suggested that acetylation of core histones, nucleosome disruption, recruitment of Pol-II and Swi2 at the *FLO1* promoter robustly induces the transcription. However, the contribution of Rlm1 in transcription of *FLO* genes has not been explored before. Our results suggest that transcriptional induction of *FLO* genes is dependent on activation of Slt2 and Rlm1. Through ChIP assay; we detected binding of Rlm1 over *FLO1* and *FLO5* (Figure 5C) promoters in flocculating *sen1-1* strain as compared to wild-type cells and *sen1-R302W*, a non-flocculating mutant. Therefore, we hypothesized that binding of Tup1 and Rlm1 at the promoter of *FLO* genes is oppositely regulated. To validate the relationship between Tup1 and Rlm1 in the regulation of *FLO* genes, we first performed ChIP assays using TBP, Pol-II, and Tup1 specific antibodies to examine occupancy at the *FLO1* and *FLO5* genes in a flocculating mutant of *Sen1* (*sen1-1*). The wild-type and non-flocculating *Sen1* mutant (*sen1-R302W*) cells were used as a control. The ChIP DNAs were amplified by using primer sets for upstream regions of *FLO1* and *FLO5* genes through RT-qPCR for analyzing the occupancy of TBP, Pol-II, and Tup1. Remarkably we found constitutive recruitment of TBP, Pol-II and a significant decrease in occupancy of Tup1 at the promoters of *FLO1* (Figure 6A) and *FLO5* (Figure 6B) in *sen1-1* cells than wild-type and non-flocculating mutant. These results suggest that signaling through the CWI pathway is critical for Rlm1 recruitment at the *FLO* genes. The low occupancy of Pol-II at the promoters of *FLO* genes in *sen1-R302W* cells could be because this substitution is known to disrupt the physical interaction between Sen1 and Rpb1 (Chinchilla *et al.* 2012). Further experiments would be required to explore the role of Rlm1, Sen1, and Tup1 in transcriptional regulation of *FLO* genes. It is quite possible that stress conditions induced by genetic or epigenetic factors trigger changes in local chromatin structure which allows Sen1 mediated binding of Rlm1 resulting in the eviction of Tup1 to induce the expression of *FLO* genes. Alternatively, the ‘loss of Sen1 function’ may prevent Tup1 to bind or recruit Rlm1 to evict Tup1. We believe this mechanism has been evolved to develop stress-tolerant phenotype.

**Figure 6:**
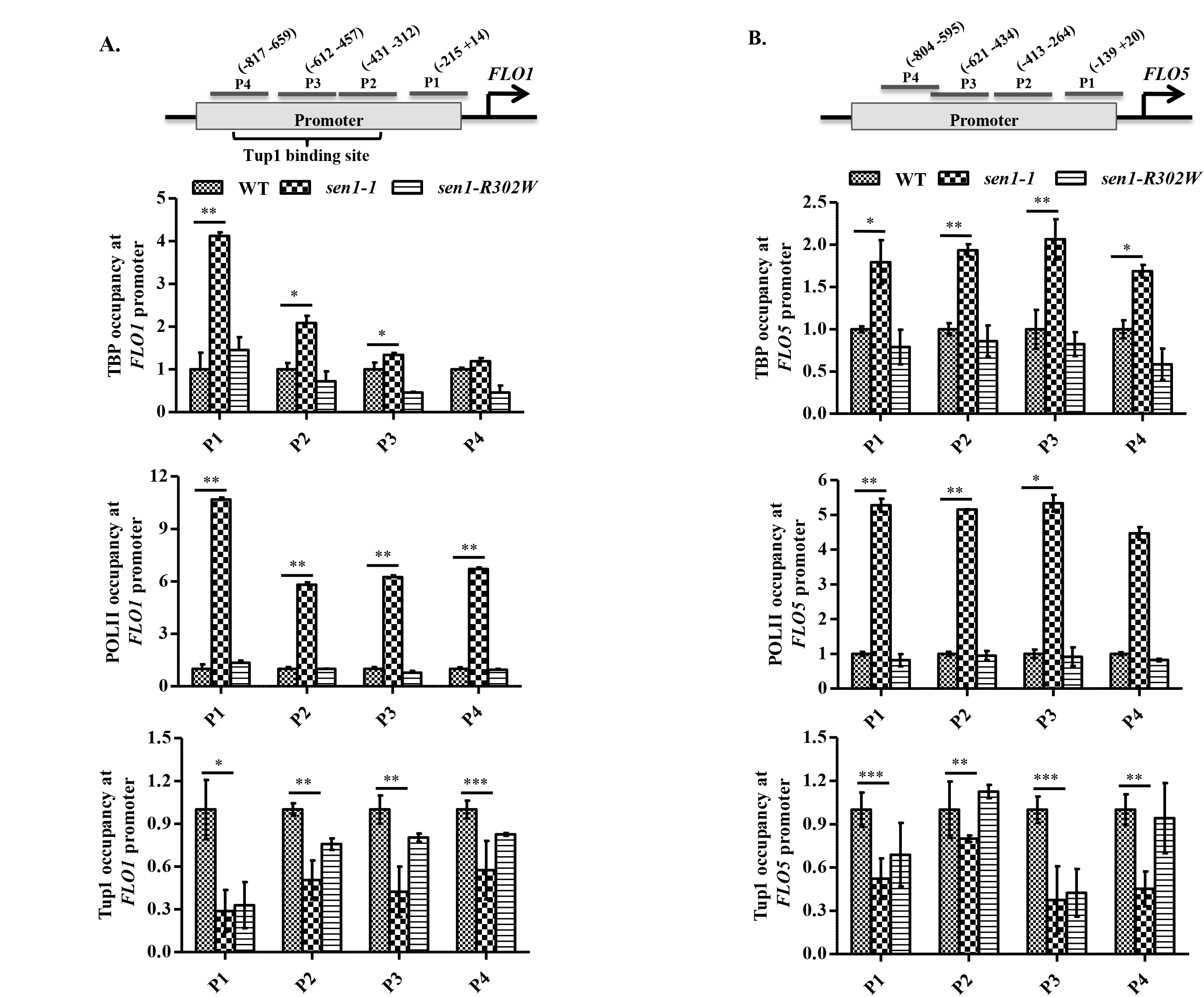
Reduced occupancy of Tup1 in flocculating mutants of *Sen1* leads to the recruitment of TBP and Pol-II at the promoter of *FLO1* and *FLO5*. ChIP assays were performed as described in materials and methods with chromatin extracts isolated from wild-type, *sen1-1*, and *sen1-R302W* cells. ChIP DNAs were analyzed by RT-qPCR. ChIP assays were performed with the following antibodies: Anti-TBP, Anti-Pol-II, and Anti-Tup1 antibodies.

A. Schematic representation indicating locations of primers of the *FLO1* gene used for amplification of ChIP DNAs is shown at the top. Chip assays were performed to check the relative occupancy of TBP, Pol-II, and Tup1 at the promoter of *FLO1*.
B. Schematic representation indicating locations of primers of the *FLO5* gene used for amplification of ChIP DNAs is shown at the top. Chip assays were performed to check the relative occupancy of TBP, Pol-II, and Tup1 at the promoter of *FLO5*. Primers of *STE6* gene were used as internal control for quantification of TBP, Pol-II and Tup1 occupancy. Bars in this data represent the difference in the fold change of relative occupancy of TBP, POLII, and Tup1 at *FLO1* and *FLO5* gene promoters in *sen1-1* and *sen1-R302W* mutants compared to that of wild-type cells. Error bars represent SD. Data shown here are the averages of three independent experiments. Statistical analysis was carried out with a two-tailed, unpaired, Student’s *t*-test to analyze differences between the *sen1-1* mutant and the wild-type cells: \**P* ≤ 0.05; \*\**P* ≤ 0.01; \*\*\**P* ≤ 0.001.

### 3.7 Slt2 is constitutively phosphorylated in flocculating histone H3 and H4 mutants leading to the Rlm1 dependent expression of *FLO* genes

Until now our results strongly suggest that the CWI pathway plays an essential role in the regulation of *FLO* genes and flocculation phenotype. However, to study whether CWI pathway dependent regulation of *FLO* genes is a general mechanism applicable to all flocculating strains or is it specific to only certain strains (for example, *Sen1* mutants) needs to be elucidated. Therefore, we decided to extend our investigations by utilizing other flocculating strains. To this end, we first looked for the additional flocculating strains. It has been earlier reported that mutants defective in COMPASS (Complex Proteins Associated with Set1) exhibit flocculation property (Dietvorst and Brandt 2008). Since subunits of COMPASS possess Histone methyltransferase activity, we decided to screen tail mutants of histone H3 and H4 for flocculation phenotype. Interestingly we were successful in finding two of the mutants; *H3R63A* and *H4R3K* with better flocculation property. First, we decided to examine the phosphorylation of Slt2 in these two mutants along with respective wild-type cells. As compared to wild-type cells, we observed more phosphorylation of Slt2 in both the mutants indicating that the CWI pathway is involved in flocculation of the histone mutants as well (Figure 7A). We then went ahead to identify the role of Rlm1 on mRNA expression of *FLO* genes, phosphorylation of Slt2, flocculation/aggregation phenotype as well as sensitivity to cell wall damaging agents in flocculating histone mutants. For this purpose, we deleted the *RLM1* gene from the mutants and wild-type cells. Relative to wild-type cells, we observed constitutive upregulation of *FLO* genes in both the flocculating histone mutants (Figure 7B and C). However, deletion of *RLM1* significantly reduced the mRNA expression of *FLO* genes, leading to a decrease in flocculation phenotype (Figure 7D and E) in both the mutants. Our careful analysis revealed low levels of Slt2 phosphorylation, reduction in expression of *FLO* genes and flocculation efficiency in the *H4R3K* mutant in comparison to *H3R63A*. Additionally, *H4R3K* cells showed less sensitivity to cell wall damaging agents relative to *H3R63A* mutant which correlates with differences in Slt2 phosphorylation (Figure S8A and S8B) (Liu *et al.* 2010). However, unlike *H3R63A*, we did not observe resistance to CR and CFW, and sensitivity to Caffeine in *H4R3K* after deletion of *RLM1*. Above results once again strongly suggest that the defect in cell wall induces the expression of *FLO* genes mediated by activation of the CWI pathway. Thus our results further suggest that the CWI pathway dependent activation of Slt2 and Rlm1 is required for the de-repression of *FLO* genes.

**Figure 7:**
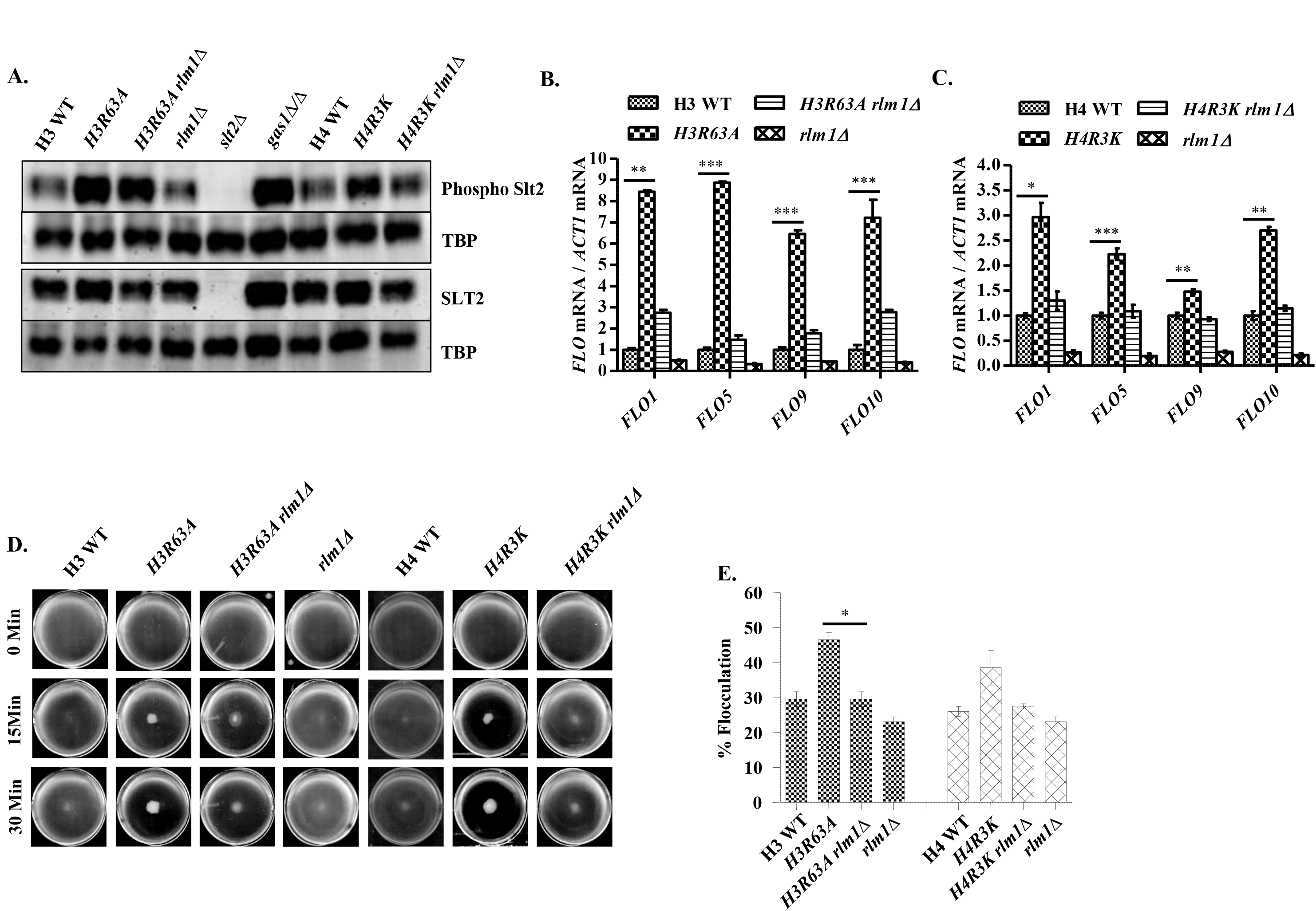
Constitutive phosphorylation of Slt2 in flocculating histone H3 and H4 mutants leads to Rlm1 dependent flocculation. A. Exponentially growing cultures of flocculating histone mutants (*H3R63A* and *H4R3K*) and respective wild-type cells with and without *RLM1* deletion were harvested; washed and whole-cell extracts were prepared by 20% TCA. Cell extracts were separated on 10% SDS-PAGE and transferred onto nitrocellulose membrane. Blots were probed with primary antibodies as indicated for checking levels of phosphorylated and non-phosphorylated forms of Slt2 by quantitative western blotting using IR Dye-labeled secondary antibodies. Protein extracts of *gas1∆/∆* and *slt2∆* strains were used as positive and negative control respectively. Blots were re-probed with anti-TBP antibody for protein loading control.
B. *B.* and C. The mRNA quantification of *FLO1, 5, 9* and *10* in flocculating histone *H3R63A* (B) and *H4R3K* (C) mutants with respective wild-type cells, with and without *RLM1* deletion by RT-qPCR. Cells were grown at 30°C, total RNAs, and cDNAs were prepared. *ACT1* mRNAs were used as internal control for quantification of *FLO* mRNAs.
C. Flocculation of flocculating histone mutants (*H3R63A* and *H4R3K*) and respective wild-type cells with and without *RLM1* deletion was recorded by taking pictures at indicated time points.
D. The % flocculation of flocculating histone mutants (*H3R63A* and *H4R3K*) and respective wild-type cells with and without *RLM1* deletion was calculated. Bars in this data represent the difference in the fold change of *FLO* gene expression (B and C) and % flocculation (E) in each mutant compared to that of wild-type cells. Error bars represent SD. Data shown here are the averages of three independent experiments. Statistical analysis was carried out with a two-tailed, unpaired, Student’s *t*-test to analyze differences between the indicated mutants and the wild-type strain (B-C) and between mutants with and without *RLM1* deletion (E):: \**P* ≤ 0.05; \*\**P* ≤ 0.01; \*\*\**P* ≤ 0.001.

### 3.8. De-repression of *FLO* genes in the absence of *TUP1* is not dependent on Slt2 and Rlm1

It is known from few studies that Cyc8-Tup1 repressor complex robustly suppresses the expression of *FLO* genes and deletion of *CYC8* or *TUP1* results in constitutive expression of *FLO* genes (Fleming *et al.* 2014). To extend our studies further, we decided to identify the role of the CWI pathway in the expression of *FLO* genes upon deletion of *TUP1* as well. To our surprise, we did not find the activation of Slt2 in strongly flocculating deletion mutants of *tup1* and *cyc8* (Figure 8A) unlike the flocculating strains of *Sen1* and Histones. Furthermore, we deleted the *SLT2* and *RLM1* in *tup1∆* mutant and measured the expression of *FLO* genes. However, we did not find any difference in the expression of *FLO* genes in *the tup1∆* mutant with and without deletion of *SLT2* or *RLM1* (Figure 8B).

**Figure 8:**
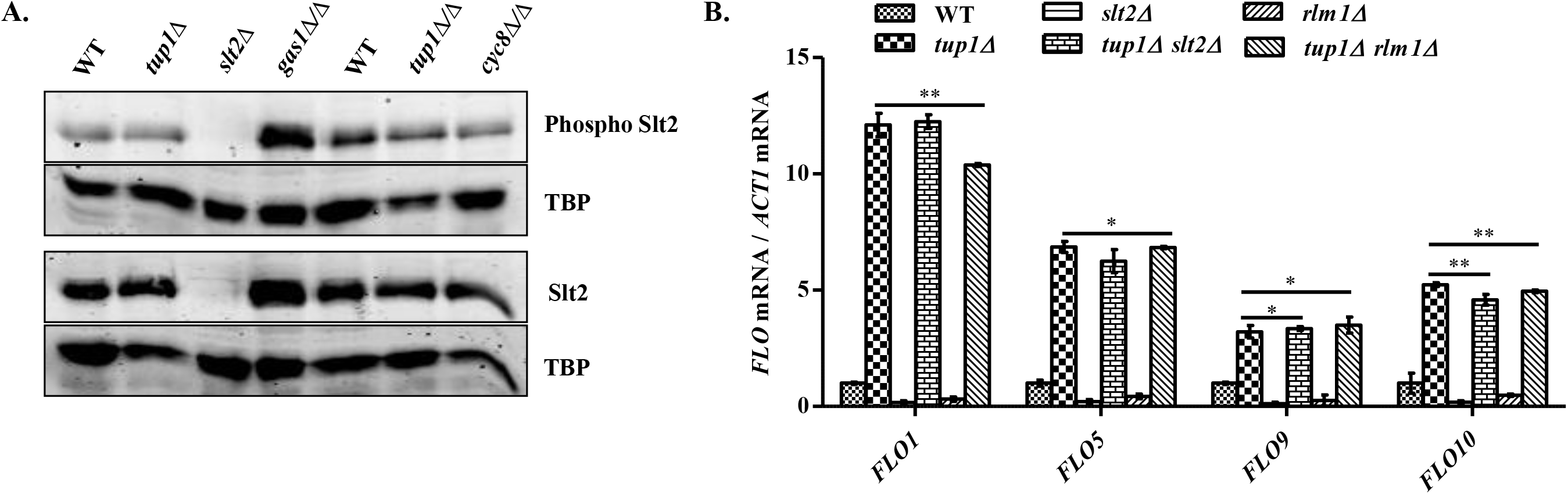
Flocculation of *tup1∆* cells is not dependent on the activation of the CWI pathway. A. Exponentially growing cultures of *tup1∆, tup1∆/∆*, and *cyc8∆/∆* with respective wild-type were harvested, washed and whole-cell extracts were prepared by 20% TCA. The cell extracts were separated on 10% SDS-PAGE and transferred onto nitrocellulose membrane. Blots were probed with primary antibodies as indicated for checking levels of phosphorylated and non-phosphorylated forms of Slt2. For protein loading control, blots were re-probed with an anti-TBP antibody. Extracts from *slt2∆* and *gas1∆/∆* cells were taken as –ve and +ve control respectively for Slt2 phosphorylation.
B. The mRNA quantification of *FLO1, 5, 9* and *10* in the *tup1∆* mutant with and without *SLT2* and *RLM1* deletion along with wild-type cells by RT-qPCR. Cells were grown at 30°C, total RNAs, and cDNAs were prepared. The *ACT1* mRNAs were used as internal control for quantification of *FLO* mRNAs. Bars in this data represent the difference in fold change of *FLO* gene expression (B) in each mutant compared to that of wild-type cells. Error bars represent SD. Data shown here are the averages of three independent experiments. Statistical analysis was carried out with a two-tailed, unpaired, Student’s *t*-test to analyze differences between the *tup1∆* mutant with double mutants, *tup1∆ slt2∆* and *tup1∆ rlm1∆* (B): \**P* ≤ 0.05; \*\**P* ≤ 0.01.

## 4. Discussion

Many factors are required for the regulation of gene expression in eukaryotes. The epigenetic and genetic factors induce structural and chemical changes to the chromatin, impacting the expression of responsive genes. Upon cell wall stress, *S. cerevisiae* activates an adaptive transcriptional response to counterbalance the stress, which is mediated by a Slt2-dependent MAPK signaling pathway. Flocculation of yeast cells is another mechanism which provides adaptation to several types of stress conditions including cell wall stress. This study was conducted to identify the role of CWI signaling pathway in the expression of *FLO* genes.

### 4.1. The yeast flocculation is regulated through a conserved MAP kinase pathway

Yeast cells are exposed to a variety of environmental stress forms and respond through a conserved MAPK pathway which induces transcription of stress-responsive genes. The specific signaling pathways through the cascade of reactions integrate into the chromatin to induce or repress the transcription of responsive genes required for survival under a variety of pathophysiological stress conditions. The environmental cell wall stress conditions result in the activation of the CWI signaling pathway leading to the upregulation of responsive genes required for maintenance of cell wall. The transcriptional program triggered by cell wall stress conditions is mediated by Slt2 and Rlm1 to induce the expression of responsive genes. Previous work has shown constitutive expression of *FLO* genes in mutants of *Sen1*, *Nab3* and *rnt1∆* cells leading to flocculation/aggregation of cells (Singh *et al.* 2015). Morphological analysis of flocculating cells (*Sen1* mutants) suggests defects in the cell wall (Figure 1C). Therefore, we hypothesized that the CWI pathway might be required for regulation of yeast flocculation. The proteins encoded by *FLO1*, *FLO5*, *FLO9*, *FLO10* and *FLO11* genes are primarily responsible for flocculation and invasive growth. The proteins encoded by *FLO* genes are glycosylphosphatidylinositol-linked glycoproteins (Dranginis *et al.* 2007). There are evidences which suggest that the mechanism of flocculation/filamentous growth is functionally conserved. First; the expression of certain human genes such as DNA methyltransferase 1 (*DNMT1*) has been reported to induce yeast flocculation (Sugiyama *et al.* 2015). Second; the expression of an activated extracellular signal-regulated kinase 1 (ERK1), one of the MAP kinases of mammalian systems in yeast has been shown to induce filamentous growth and cell wall remodeling (Atienza *et al.* 2000). Third; most of the yeast cell wall flocculins and human cell membrane adhesins consists of amyloid-forming amino acid sequences (Rameau *et al.* 2016; Ramsook *et al.* 2010). Fourth, certain mutations in conserved histone proteins and deletion of a histone-modifying enzyme (*swd3∆*) also induce flocculation of yeast cells (Dietvorst and Brandt 2008). Mutations in histones and histone-modifying enzymes have been implicated in diseases including cancer (Shilatifard 2012). Fifth, during current studies we detected constitutive activation of a conserved MAP kinase (Slt2) dependent pathway known as Cell wall integrity (CWI) in flocculating cells (Figure 2B, Figure S4A and Figure 7A). It is quite possible that under certain stress conditions due to epigenetic or genetic changes, human cell membrane adhesins like yeast cell wall flocculins may induce morphological changes affecting cell-cell communication and pathological properties. Many cellular activities such as proliferation, differentiation, and cell death have been shown to be controlled by signaling pathways involving specific MAP kinases (Kim and Choi 2010). The defects in MAP kinases are implicated in many human diseases including Alzheimer’s disease, Parkinson’s disease, amyotrophic lateral sclerosis (ALS) and cancers (Kim and Choi 2010). Many mutations in human senataxin (yeast Sen1) have been correlated with neurological disorders, AOA2 and ALS4. However, the underlying mechanism for AOA2 and ALS4 as well as induction of flocculation due to mutations in yeast Sen1 is not clear. Our observations suggest that dysregulation in MAP kinase pathway might be responsible for AOA2 and ALS4 disorders associated with mutations in human senataxin.

### 4.2. The Rlm1 is required for the de-repression of *FLO* genes under stress conditions

To dissect the molecular mechanism of yeast flocculation, we performed a variety of experiments. First, we found that flocculating strains exhibit enlarged cell size, higher chitin content and rough cell surface morphology which is indicative of alterations in the cell wall. We then decided to examine the CWI pathway by testing the phosphorylation of Slt2, one of the key MAP kinases. Interestingly, we detected constitutive phosphorylation of Slt2 in flocculating strains, not only in *Sen1* mutants but also in flocculating histone mutant cells, suggesting that the CWI signaling pathway indeed regulates the de-repression of *FLO* genes. It is known from several studies that phosphorylated form of Slt2 enters into the nucleus and which acts as a kinase to activate the Rlm1, a transcription factor which in turn induces the transcription of cell wall damage responsive genes by physically interacting with SWI/SNF chromatin remodeling complex and recruitment of general transcription factors. However, the significance of Rlm1 in de-repression of *FLO* genes has not been explored before. Excitingly, in addition to constitutive activation of Slt2, we also found constitutive recruitment of Pol-II and TBP, and reduced occupancy of Tup1 at the promoters of *FLO* genes in flocculating cells revealing the role of CWI signaling pathway in de-repression of *FLO* genes. Furthermore, we found reduced expression of *FLO* genes and suppression of flocculation phenotype upon deletion of *SLT2* or *RLM1* (Figure 4 and 5). To get more insight, we also measured the basal expression of *FLO* genes in deletion mutant cells of the CWI pathway. Remarkably we found a reduction in basal expression of *FLO* genes in all the deletion mutants of the CWI pathway except *mlp1∆* cells. Since basal expression of *FLO* genes in *mlp1∆* mutant did not decline, unlike other mutants, suggesting that expression of *FLO* genes is exclusively regulated through Rlm1 branch of CWI signaling pathway. These observations motivated us to hypothesize that Rlm1 is required for the transcription of *FLO* genes by binding to the promoters of *FLO* genes under stress conditions. We, therefore, searched for the Rlm1 binding site at the *FLO* genes by using a previously identified DNA sequences, TA(A/T)_4_TAG as Rlm1 binding sequences. We found that Rlm1 binding sites are indeed available at the upstream regions of *FLO1, 5, 9* and *10* genes, overlapping with the Tup1 binding sites which were confirmed by ChIP assay (Figure 5A, B, C). Furthermore, to examine whether phosphorylation of Slt2 has any correlation or not with transcriptional induction of *FLO* genes during cell wall stress, we artificially induced cell wall damage condition by exposing wild-type cells to CFW and higher temperature. As per our assumption, in both the conditions, we observed an increase in phosphorylation of Slt2 and upregulation of *FLO1, 5*, and *9* leading to flocculation of cells. These results indicate the stress-dependent role of the CWI pathway in transcriptional de-repression of *FLO* genes via activation of Slt2 and Rlm1. Furthermore, to confirm the role of the CWI pathway in the regulation of *FLO* genes, first, we performed similar experiments by using additional flocculating strains other than *Sen1* mutants. Through the screening of histone mutants, we found few mutants with the flocculating property. Out of these, we selected two mutants which showed better flocculation phenotype. We detected an increase in phosphorylation of Slt2 and induction of *FLO* genes in flocculating histone mutants as well, which further suggest that activation of Slt2p and Rlm1p-dependent CWI pathway is the general mechanism required for flocculation of yeast cells. Next, we also tested phosphorylation of Slt2 in *CYC8* and *TUP1* deleted cells which are known to possess strong flocculation property (Fleming *et al.* 2014). To our surprise, we did not detect any considerable increase in phosphorylation of Slt2 in flocculating *CYC8* and *TUP1* deleted cells. Moreover, upon deletion of *SLT2* or *RLM1* in *tup1∆* cells also did not affect the expression of *FLO* genes. Future investigations would be required to understand the mechanism for Slt2-independent induction of *FLO* genes in *tup1∆* and *cyc8∆* cells.

We hypothesize two possibilities for Slt2-independent induction of *FLO* genes in the absence of *CYC8* and *TUP1*; one, Rlm1-dependent and second, Rlm1-independent. Based on Rlm1-dependent mechanism we propose that in the absence of *CYC8* or *TUP1*, the basal level of Rlm1p is sufficient to bind at the freely accessible Rlm1 binding sites at the promoter of *FLO* genes. Tup1p repressor in an unstressed condition probably occupies the Rlm1 binding site as binding sites for these two factors are overlapping. According to the second Rlm1-independent mechanism, possibly the nucleosome positioning is disrupted due to the opportunistic recruitment of SWI/SNF complex in the absence of Tup1p leading to the formation of PIC causing robust transcriptional de-repression of *FLO* genes. However, to support or refute these mechanisms further investigations are needed. It has been shown earlier that Cyc8-Tup1 repressor complex regulates the chromatin structure and gene expression of repressed genes by influencing Isw2-dependent nucleosome positioning (Rizzo *et al.* 2011; Zhang and Reese 2004) which supports our conclusions to some extent. Despite many studies, the precise mechanism for the regulation of *FLO* genes under cell wall stress conditions has not been explored before. Our results suggest that under cell wall stress conditions, activated Slt2 triggers the recruitment of active Rlm1 results in eviction of Tup1, recruitment of TBP and Pol-II, to induce the expression of *FLO* genes (Figure 9). Several studies have established the fact that nucleosome positioning influences the accessibility of the transcription factors binding sites at the promoters. The positioning of nucleosomes in yeast is primarily determined by two major chromatin remodeling complexes; ISW2 and SWI2/SNF2 (Tomar *et al.* 2009). Our ChIP assays suggest that Rlm1 is constitutively bound to the promoter of *FLO* genes in flocculating cells probably due to disruption of nucleosome positioning. The occupancy of Tup1 in the absence of Rlm1 perhaps maintains the repressed state of *FLO* genes due to nucleosome positioning. Further *in vivo* and *in vitro* investigations would be required to establish a precise relationship between Rlm1, Tup1 and chromatin remodeling.

**Figure 9:**
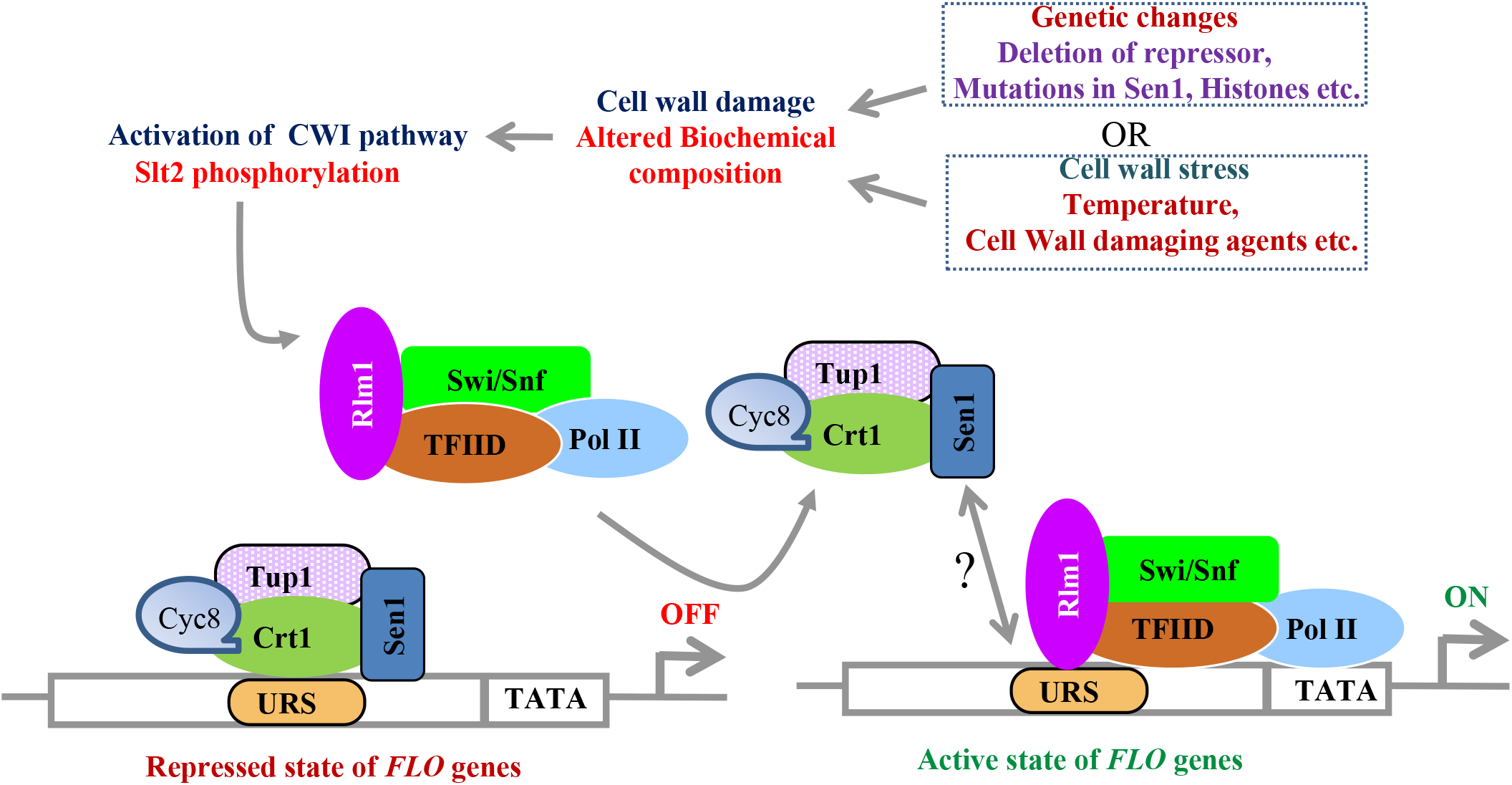
Model illustrating the role of the CWI pathway in the regulation of *FLO* genes through Sen1 mediated antagonism between Rlm1 and Tup1. Genetic (mutations in genes) and stress conditions (cell wall damaging agents and higher temperature) alters the biochemical composition of yeast cell wall results in activation of the CWI signaling pathway via phosphorylation of Slt2. Activated Slt2 enters into the nucleus and activates the Rlm1. Activated Rlm1 along with general transcription factors; TBP, Pol-II, and SWI/SNF is recruited to form Pre Initiation Complex (PIC) and dissociates the Tup1 repressor, converting repressed (OFF) state of *FLO* genes to transcriptional active (ON) state. To induce the expression of *FLO* genes, Sen1 probably cooperates with multiple proteins including Rlm1, factors of general transcription machinery leading to the eviction of Tup1 repressor.

## 5. Conclusion

The novel Rlm1-dependent mechanism for the transcription of *FLO* genes has certainly enhanced our knowledge about the molecular biology of yeast flocculation/biofilm formation. In summary, our studies are expected to serve two primary purposes, first; strengthening the understanding about yeast biofilm biology which may help us to tackle the antifungal resistance and therapies for biofilm-based fungal diseases and second; in biotechnological applications as flocculating yeast cells are useful for fermentation processes to develop antimicrobials and biofuel.

## Author contribution

RST, SKS, VS conceived and designed the study. SKS and RK performed the experiments. Results were analyzed by RST, SKS, RK, and VS. RST, SKS, RK, and VS wrote the manuscript. SKS, RK, and VS prepared all the figures. All authors reviewed the results and approved the final version of the manuscript.

## Acknowledgments

We thank Michael Culbertson (University of Wisconsin), Jeffry Corden (Johns Hopkins University School of Medicine), Steve Buratowski (Harvard Medical School), Daniel Reines (Emory University), Anita H. Corbet (Emory University), Sherif Abou Elela (University of Sherbrooke) and David E. Levin (Boston University) for providing us the Sen1, Nrd1, Nab3, TRAMP, Rnt1 and CWI pathway mutant strains of yeast respectively. We acknowledge Joseph Reese (Pennsylvania State University) for the generous gift of TBP and Tup1 antibodies. Council of Scientific and Industrial Research (CSIR), India is acknowledged for fellowship support to SKS and RK. Members of the chromatin biology laboratory are recognized for their advice and helpful discussions throughout this work.

## References

Amberg, D. C., D. J. Burke, D. Burke, J. N. Strathern and C. S. H. Laboratory, 2005 Methods in Yeast Genetics: A Cold Spring Harbor Laboratory Course Manual. Cold Spring Harbor Laboratory Press.

Arroyo, J., V. Farkas, A. B. Sanz and E. Cabib, 2016 ‘Strengthening the fungal cell wall through chitin-glucan cross-links: effects on morphogenesis and cell integrity. Cell Microbiol 18: 1239–1250.

Atienza, J. M., M. Suh, I. Xenarios, R. Landgraf and J. Colicelli, 2000 Human ERK1 induces filamentous growth and cell wall remodeling pathways in Saccharomyces cerevisiae. J Biol Chem 275: 20638–20646.

Baccarelli, A., and V. Bollati, 2009 Epigenetics and environmental chemicals. Curr Opin Pediatr 21: 243–251.

Barua, S., L. Li, P. N. Lipke and A. M. Dranginis, 2016 Molecular Basis for Strain Variation in the Saccharomyces cerevisiae Adhesin Flo11p. mSphere 1.

Bermejo, C., E. Rodriguez, R. Garcia, J. M. Rodriguez-pena, M. L. Rodriguez De La Concepcion et al., 2008 The sequential activation of the yeast HOG and SLT2 pathways is required for cell survival to cell wall stress. Mol Biol Cell 19: 1113–1124.

Bester, M. C., D. Jacobson and F. F. Bauer, 2012 Many Saccharomyces cerevisiae Cell Wall Protein Encoding Genes Are Coregulated by Mss11, but Cellular Adhesion Phenotypes Appear Only Flo Protein Dependent. G3 (Bethesda) 2: 131–141.

Birkaya, B., A. Maddi, J. Joshi, S. J. Free and P. J. Cullen, 2009 Role of the cell wall integrity and filamentous growth mitogen-activated protein kinase pathways in cell wall remodeling during filamentous growth. Eukaryot Cell 8: 1118–1133.

Bony, M., P. Barre and B. Blondin, 1998 Distribution of the flocculation protein, flop, at the cell surface during yeast growth: the availability of flop determines the flocculation level. Yeast 14: 25–35.

Bou Zeidan, M., G. Zara, C. Viti, F. Decorosi, I. Mannazzu et al., 2014 L-histidine inhibits biofilm formation and FLO11-associated phenotypes in Saccharomyces cerevisiae flor yeasts. PLoS One 9: e112141.

Chandra, J., D. M. Kuhn, P. K. Mukherjee, L. L. Hoyer, T. Mccormick et al., 2001 Biofilm formation by the fungal pathogen Candida albicans: development, architecture, and drug resistance. J Bacteriol 183: 5385–5394.

Chavel, C. A., L. M. Caccamise, B. Li and P. J. Cullen, 2014 Global regulation of a differentiation MAPK pathway in yeast. Genetics 198: 1309–1328.

Chavel, C. A., H. M. Dionne, B. Birkaya, J. Joshi and P. J. Cullen, 2010 Multiple signals converge on a differentiation MAPK pathway. PLoS Genet 6: e1000883.

Chen, R. E., and J. Thorner, 2007 Function and regulation in MAPK signaling pathways: lessons learned from the yeast Saccharomyces cerevisiae. Biochim Biophys Acta 1773: 1311–1340.

Chen, X., K. Poorey, M. N. Carver, U. Muller, S. Bekiranov*, et al.*, 2017 Transcriptomes of six mutants in the Sen1 pathway reveal combinatorial control of transcription termination across the Saccharomyces cerevisiae genome. PLoS Genet 13: e1006863.

Chinchilla, K., J. B. Rodriguez-molina, D. Ursic, J. S. Finkel, A. Z. Ansari et al., 2012 Interactions of Sen1, Nrd1, and Nab3 with multiple phosphorylated forms of the Rpb1 C-terminal domain in Saccharomyces cerevisiae. Eukaryot Cell 11: 417–429.

Church, M., K. C. Smith, M. M. Alhussain, S. Pennings and A. B. Fleming, 2017 Sas3 and Ada2(Gcn5)-dependent histone H3 acetylation is required for transcription elongation at the de-repressed FLO1 gene. Nucleic Acids Res 45: 4413–4430.

Cohen, S., N. Puget, Y. L. Lin, T. Clouaire, M. Aguirrebengoa et al., 2018 Senataxin resolves RNA:DNA hybrids forming at DNA double-strand breaks to prevent translocations. Nat Commun 9: 533.

De Nobel, H., C. Ruiz, H. Martin, W. Morris, S. Brul et al., 2000 Cell wall perturbation in yeast results in dual phosphorylation of the Slt2/Mpk1 MAP kinase and in an Slt2-mediated increase in FKS2-lacZ expression, glucanase resistance and thermotolerance. Microbiology 146 (Pt 9): 2121–2132.

Degreif, D., T. De Rond, A. Bertl, J. D. Keasling and I. Budin, 2017 Lipid engineering reveals regulatory roles for membrane fluidity in yeast flocculation and oxygen-limited growth. Metab Eng 41: 46–56.

Dietvorst, J., and A. Brandt, 2008 Flocculation in Saccharomyces cerevisiae is repressed by the COMPASS methylation complex during high-gravity fermentation. Yeast 25: 891–901.

Dodou, E., and R. Treisman, 1997 The Saccharomyces cerevisiae MADS-box transcription factor Rlm1 is a target for the Mpk1 mitogen-activated protein kinase pathway. Mol Cell Biol 17: 1848–1859.

Dranginis, A. M., J. M. Rauceo, J. E. Coronado and P. N. Lipke, 2007 A biochemical guide to yeast adhesins: glycoproteins for social and antisocial occasions. Microbiol Mol Biol Rev 71: 282–294.

Ene, I. V., L. A. Walker, M. Schiavone, K. K. Lee, H. Martin-yken et al., 2015 Cell Wall Remodeling Enzymes Modulate Fungal Cell Wall Elasticity and Osmotic Stress Resistance. MBio 6: e00986.

Fasken, M. B., R. N. Laribee and A. H. Corbett, 2015 Nab3 facilitates the function of the TRAMP complex in RNA processing via recruitment of Rrp6 independent of Nrd1. PLoS Genet 11: e1005044.

Fichtner, L., F. Schulze and G. H. Braus, 2007 Differential Flo8p-dependent regulation of FLO1 and FLO11 for cell-cell and cell-substrate adherence of S. cerevisiae S288c. Mol Microbiol 66: 1276–1289.

Finkel, J. S., K. Chinchilla, D. Ursic and M. R. Culbertson, 2010 Sen1p performs two genetically separable functions in transcription and processing of U5 small nuclear RNA in Saccharomyces cerevisiae. Genetics 184: 107–118.

Fleming, A. B., S. Beggs, M. Church, Y. Tsukihashi and S. Pennings, 2014 The yeast Cyc8-Tup1 complex cooperates with Hda1p and Rpd3p histone deacetylases to robustly repress transcription of the subtelomeric FLO1 gene. Biochim Biophys Acta 1839: 1242–1255.

Fleming, A. B., and S. Pennings, 2001 Antagonistic remodelling by Swi-Snf and Tup1-Ssn6 of an extensive chromatin region forms the background for FLO1 gene regulation. EMBO J 20: 5219–5231.

Fox, M. J., H. Gao, W. R. Smith-kinnaman, Y. Liu and A. L. Mosley, 2015 The exosome component Rrp6 is required for RNA polymerase II termination at specific targets of the Nrd1-Nab3 pathway. PLoS Genet 11: e1004999.

Garcia, R., C. Bermejo, C. Grau, R. Perez, J. M. Rodriguez-pena et al., 2004 The global transcriptional response to transient cell wall damage in Saccharomyces cerevisiae and its regulation by the cell integrity signaling pathway. J Biol Chem 279: 15183–15195.

Garcia, R., J. M. Rodriguez-pena, C. Bermejo, C. Nombela and J. Arroyo, 2009 The high osmotic response and cell wall integrity pathways cooperate to regulate transcriptional responses to zymolyase-induced cell wall stress in Saccharomyces cerevisiae. J Biol Chem 284: 10901–10911.

Garcia, R., A. B. Sanz, J. M. Rodriguez-pena, C. Nombela and J. Arroyo, 2016 Rlm1 mediates positive autoregulatory transcriptional feedback that is essential for Slt2-dependent gene expression. J Cell Sci 129: 1649–1660.

Golla, U., V. Singh, G. K. Azad, P. Singh, N. Verma et al., 2013 Sen1p contributes to genomic integrity by regulating expression of ribonucleotide reductase 1 (RNR1) in Saccharomyces cerevisiae. PLoS One 8: e64798.

Goossens, K. V., F. S. Ielasi, I. Nookaew, I. Stals, L. Alonso-sarduy et al., 2015 Molecular mechanism of flocculation self-recognition in yeast and its role in mating and survival. MBio 6.

Goossens, K. V., C. Stassen, I. Stals, D. S. Donohue, B. Devreese et al., 2011 The N-terminal domain of the Flo1 flocculation protein from Saccharomyces cerevisiae binds specifically to mannose carbohydrates. Eukaryot Cell 10: 110–117.

Groh, M., L. O. Albulescu, A. Cristini and N. Gromak, 2017 Senataxin: Genome Guardian at the Interface of Transcription and Neurodegeneration. J Mol Biol 429: 3181–3195.

Grunseich, C., I. X. Wang, J. A. Watts, J. T. Burdick, R. D. Guber et al., 2018 Senataxin Mutation Reveals How R-Loops Promote Transcription by Blocking DNA Methylation at Gene Promoters. Mol Cell 69: 426–437 e427.

Guo, B., C. A. Styles, Q. Feng and G. R. Fink, 2000 A Saccharomyces gene family involved in invasive growth, cell-cell adhesion, and mating. Proc Natl Acad Sci U S A 97: 12158–12163.

Gustin, M. C., J. Albertyn, M. Alexander and K. Davenport, 1998 MAP kinase pathways in the yeast Saccharomyces cerevisiae. Microbiol Mol Biol Rev 62: 1264–1300.

Hope, E. A., C. J. Amorosi, A. W. Miller, K. Dang, C. S. Heil et al., 2017 Experimental Evolution Reveals Favored Adaptive Routes to Cell Aggregation in Yeast. Genetics 206: 1153–1167.

Janke, C., M. M. Magiera, N. Rathfelder, C. Taxis, S. Reber et al., 2004 A versatile toolbox for PCR-based tagging of yeast genes: new fluorescent proteins, more markers and promoter substitution cassettes. Yeast 21: 947–962.

Jimenez-sanchez, M., V. J. Cid and M. Molina, 2007 Retrophosphorylation of Mkk1 and Mkk2 MAPKKs by the Slt2 MAPK in the yeast cell integrity pathway. J Biol Chem 282: 31174–31185.

Jung, U. S., and D. E. Levin, 1999 Genome-wide analysis of gene expression regulated by the yeast cell wall integrity signalling pathway. Mol Microbiol 34: 1049–1057.

Kim, E. K., and E. J. Choi, 2010 Pathological roles of MAPK signaling pathways in human diseases. Biochim Biophys Acta 1802: 396–405.

Kim, J., and M. D. Rose, 2015 Stable Pseudohyphal Growth in Budding Yeast Induced by Synergism between Septin Defects and Altered MAP-kinase Signaling. PLoS Genet 11: e1005684.

Kim, K. Y., and D. E. Levin, 2011 Mpk1 MAPK association with the Paf1 complex blocks Sen1-mediated premature transcription termination. Cell 144: 745–756.

Kim, K. Y., A. W. Truman, S. Caesar, G. Schlenstedt and D. E. Levin, 2010 Yeast Mpk1 cell wall integrity mitogen-activated protein kinase regulates nucleocytoplasmic shuttling of the Swi6 transcriptional regulator. Mol Biol Cell 21: 1609–1619.

Lee, T. I., and R. A. Young, 2013 Transcriptional regulation and its misregulation in disease. Cell 152: 1237–1251.

Levin, D. E., 2005 Cell wall integrity signaling in Saccharomyces cerevisiae. Microbiol Mol Biol Rev 69: 262–291.

Levin, D. E., 2011 Regulation of cell wall biogenesis in Saccharomyces cerevisiae: the cell wall integrity signaling pathway. Genetics 189: 1145–1175.

Lipke, P. N., M. C. Garcia, D. Alsteens, C. B. Ramsook, S. A. Klotz et al., 2012 Strengthening relationships: amyloids create adhesion nanodomains in yeasts. Trends Microbiol 20: 59–65.

Lipke, P. N., and C. Hull-pillsbury, 1984 Flocculation of Saccharomyces cerevisiae tup1 mutants. J Bacteriol 159: 797–799.

Liu, X., X. Zhang and Z. Zhang, 2010 Cu,Zn-superoxide dismutase is required for cell wall structure and for tolerance to cell wall-perturbing agents in Saccharomyces cerevisiae. FEBS Lett 584: 1245–1250.

Liu, Z., R. Li, Q. Dong, L. Bian, X. Li et al., 2015 Characterization of the non-sexual flocculation of fission yeast cells that results from the deletion of ribosomal protein L32. Yeast 32: 439–449.

Livak, K. J., and T. D. Schmittgen, 2001 Analysis of relative gene expression data using real-time quantitative PCR and the 2(-Delta Delta C(T)) Method. Methods 25: 402–408.

Madden, K., Y. J. Sheu, K. Baetz, B. Andrews and M. Snyder, 1997 SBF cell cycle regulator as a target of the yeast PKC-MAP kinase pathway. Science 275: 1781–1784.

Martin-tumasz, S., and D. A., Brow, 2015 Saccharomyces cerevisiae Sen1 Helicase Domain Exhibits 5’- to 3’-Helicase Activity with a Preference for Translocation on DNA Rather than RNA. J Biol Chem 290: 22880–22889.

Mcmurray, C. T., and J. A. Tainer, 2003 Cancer, cadmium and genome integrity. Nat Genet 34: 239–241.

O’Connor, M. J., 2015 Targeting the DNA Damage Response in Cancer. Mol Cell 60: 547–560.

Pujol-carrion, N., M. I. Petkova, L. Serrano and M. A. De La Torre-ruiz, 2013 The MAP kinase Slt2 is involved in vacuolar function and actin remodeling in Saccharomyces cerevisiae mutants affected by endogenous oxidative stress. Appl Environ Microbiol 79: 6459–6471.

Rameau, R. D., D. N. Jackson, A. Beaussart, Y. F. Dufrene and P. N. Lipke, 2016 The Human Disease-Associated Abeta Amyloid Core Sequence Forms Functional Amyloids in a Fungal Adhesin. MBio 7: e01815–01815.

Ramsook, C. B., C. Tan, M. C. Garcia, R. Fung, G. Soybelman et al., 2010 Yeast cell adhesion molecules have functional amyloid-forming sequences. Eukaryot Cell 9: 393–404.

Rizzo, J. M., P. A. Mieczkowski and M. J. Buck, 2011 Tup1 stabilizes promoter nucleosome positioning and occupancy at transcriptionally plastic genes. Nucleic Acids Res 39: 8803–8819.

Saha, S., A. A. Anilkumar and S. Mayor, 2016 GPI-anchored protein organization and dynamics at the cell surface. J Lipid Res 57: 159–175.

Sanz, A. B., R. Garcia, J. M. Rodriguez-pena, S. Diez-muniz, C. Nombela et al., 2012 Chromatin remodeling by the SWI/SNF complex is essential for transcription mediated by the yeast cell wall integrity MAPK pathway. Mol Biol Cell 23: 2805–2817.

Sanz, A. B., R. Garcia, J. M. Rodriguez-pena, C. Nombela and J. Arroyo, 2016 Cooperation between SAGA and SWI/SNF complexes is required for efficient transcriptional responses regulated by the yeast MAPK Slt2. Nucleic Acids Res 44: 7159–7172.

Sariki, S. K., P. K. Sahu, U. Golla, V. Singh, G. K. Azad et al., 2016 Sen1, the homolog of human Senataxin, is critical for cell survival through regulation of redox homeostasis, mitochondrial function, and the TOR pathway in Saccharomyces cerevisiae. FEBS J 283: 4056–4083.

Schmitt, M. E., T. A. Brown and B. L. Trumpower, 1990 A rapid and simple method for preparation of RNA from Saccharomyces cerevisiae. Nucleic Acids Res 18: 3091–3092.

Scrimale, T., L. Didone, K. L. De Mesy Bentley and D. J. Krysan, 2009 The unfolded protein response is induced by the cell wall integrity mitogen-activated protein kinase signaling cascade and is required for cell wall integrity in Saccharomyces cerevisiae. Mol Biol Cell 20: 164–175.

Shilatifard, A., 2012 The COMPASS family of histone H3K4 methylases: mechanisms of regulation in development and disease pathogenesis. Annu Rev Biochem 81: 65–95.

Sim, L., M. Groes, K. Olesen and A. Henriksen, 2013 Structural and biochemical characterization of the N-terminal domain of flocculin Lg-Flo1p from Saccharomyces pastorianus reveals a unique specificity for phosphorylated mannose. FEBS J 280: 1073–1083.

Singh, V., G. K. Azad, S. K. Sariki and R. S. Tomar, 2015 Flocculation in Saccharomyces cerevisiae is regulated by RNA/DNA helicase Sen1p. FEBS Lett 589: 3165–3174.

Smith, E., and A. Shilatifard, 2010 The chromatin signaling pathway: diverse mechanisms of recruitment of histone-modifying enzymes and varied biological outcomes. Mol Cell 40: 689–701.

Smith, K. T., and J. L. Workman, 2012 Chromatin proteins: key responders to stress. PLoS Biol 10: e1001371.

Smith, R. L., and A. D. Johnson, 2000 Turning genes off by Ssn6-Tup1: a conserved system of transcriptional repression in eukaryotes. Trends Biochem Sci 25: 325–330.

Smukalla, S., M. Caldara, N. Pochet, A. Beauvais, S. Guadagnini et al., 2008 FLO1 is a variable green beard gene that drives biofilm-like cooperation in budding yeast. Cell 135: 726–737.

Soares, E. V., 2011 Flocculation in Saccharomyces cerevisiae: a review. J Appl Microbiol 110: 1–18.

Sugiyama, K., M. Takamune, H. Furusawa and M. Honma, 2015 Human DNA methyltransferase gene-transformed yeasts display an inducible flocculation inhibited by 5-aza-2’-deoxycytidine. Biochem Biophys Res Commun 456: 689–694.

Tang, S. Y., W. Zhang, R. Soffe, S. Nahavandi, R. Shukla et al., 2014 High resolution scanning electron microscopy of cells using dielectrophoresis. PLoS One 9: e104109.

Taymaz-nikerel, H., A. Cankorur-cetinkaya and B. Kirdar, 2016 Genome-Wide Transcriptional Response of Saccharomyces cerevisiae to Stress-Induced Perturbations. Front Bioeng Biotechnol 4: 17.

Thorsen, M., Y. Di, C. Tangemo, M. Morillas, D. Ahmadpour et al., 2006 The MAPK Hog1p modulates Fps1p-dependent arsenite uptake and tolerance in yeast. Mol Biol Cell 17: 4400–4410.

Tomar, R. S., J. N. Psathas, H. Zhang, Z. Zhang and J. C. Reese, 2009 A novel mechanism of antagonism between ATP-dependent chromatin remodeling complexes regulates RNR3 expression. Mol Cell Biol 29: 3255–3265.

Tomar, R. S., S. Zheng, D. Brunke-reese, H. N. Wolcott and J. C. Reese, 2008 Yeast Rap1 contributes to genomic integrity by activating DNA damage repair genes. EMBO J 27: 1575–1584.

Ursic, D., K. Chinchilla, J. S. Finkel and M. R. Culbertson, 2004 Multiple protein/protein and protein/RNA interactions suggest roles for yeast DNA/RNA helicase Sen1p in transcription, transcription-coupled DNA repair and RNA processing. Nucleic Acids Res 32: 2441–2452.

Vasiljeva, L., M. Kim, H. Mutschler, S. Buratowski and A. Meinhart, 2008 The Nrd1-Nab3-Sen1 termination complex interacts with the Ser5-phosphorylated RNA polymerase II C-terminal domain. Nat Struct Mol Biol 15: 795–804.

Verstrepen, K. J., and F. M. Klis, 2006 Flocculation, adhesion and biofilm formation in yeasts. Mol Microbiol 60: 5–15.

Vihervaara, A., F. M. Duarte and J. T. Lis, 2018 Molecular mechanisms driving transcriptional stress responses. Nat Rev Genet.

Wach, A., 1996 PCR-synthesis of marker cassettes with long flanking homology regions for gene disruptions in S. cerevisiae. Yeast 12: 259–265.

Zhang, Z., and J. C. Reese, 2004 Ssn6-Tup1 requires the ISW2 complex to position nucleosomes in Saccharomyces cerevisiae. EMBO J 23: 2246–2257.

